# Long-Run Olfactory Enrichment Promotes Non-Olfactory Cognition, Noradrenergic Plasticity And Remodeling Of Brain Functional Connectivity In Aged Mice

**DOI:** 10.1101/2023.04.28.538433

**Authors:** Claire Terrier, Juliette Greco-Vuilloud, Matthias Cavelius, Marc Thévenet, Nathalie Mandairon, Anne Didier, Marion Richard

## Abstract

Normal brain aging is accompanied by functional and structural changes, leading to cognitive decline. A high level of cognitive stimulation during life is associated with improved cognitive performances in elderly, forming the so-called cognitive reserve whose cellular mechanisms remain largely unknown. Noradrenaline has been proposed as a molecular link between environmental stimulation and the constitution of the cognitive reserve. Taking advantage of the ability of olfactory stimulation to trigger noradrenaline release, we used repeated olfactory enrichment sessions distributed over the mouse lifespan to enable the cognitive reserve buildup. Mice submitted to lifelong olfactory enrichment whether started in early or late adulthood, displayed improved olfactory discrimination at late ages. Interestingly, benefits extended to spatial memory and cognitive flexibility and olfactory and non-olfactory cognitive performances correlated with increased noradrenergic innervation in the olfactory bulb and dorsal hippocampus. Finally, using c-Fos mapping and connectivity analysis, we reported task-specific remodeling of functional neural networks in aged mice with increased network specialization or enlargement in an olfactory discrimination or a spatial memory task respectively. We thus propose long-run olfactory enrichment as a mouse model of the cognitive reserve buildup relying on plasticity of the noradrenergic system and network remodeling to promote better cognitive aging.

## INTRODUCTION

Aging is an inevitable and complex biological phenomenon associated with a progressive decline of sensory, motor and cognitive functions with time, affecting quality of life and health. Apart from any age-related neurodegenerative diseases, brain aging is accompanied by functional and structural changes, including brain networks’ remodeling (Damoiseaux, 2017) and alterations in synaptic function (van der Zee, 2015). A potentially powerful tool to achieve successful brain aging is to boost the cognitive reserve. This concept was initially proposed to explain the discrepancy between considerable histological brain alterations and relatively preserved cognitive performances in some Alzheimer’s patients and has been extended to normal aging (Grady, 2012; Barulli and Stern, 2013; Reuter-Lorenz and Park, 2014; Cabeza et al., 2018; Stern et al., 2019). Epidemiological studies indicate that a high level of brain stimulation through education, occupational attainment or social activities, is associated with changes in neural activity and connectivity possibly supporting better capacity or adaptability of neural networks in humans. These observations prompted the development of training strategies in the auditory system, with positive outcomes on performances and remodeling of brain activation in aged subjects (*e.g.* Holman and de Villers-Sidani, 2014).

Despite a growing interest toward the cognitive reserve, the mechanisms underlying the implementation of various experiences into brain networks to allow better performances during the cognitive reserve buildup remain mostly unknown. The noradrenergic theory of the cognitive reserve proposes the neuromodulator noradrenaline (NA) as a key player in this process (Robertson, 2013; Clewett et al., 2016; Mather and Harley, 2016). This hypothesis arose from several clinical and experimental observations. The *Locus Coeruleus* (the main source of cortical NA) is vulnerable to aging and its anatomical integrity is positively associated with preserved cognitive functions in healthy humans (Wilson et al., 2013; Clewett et al., 2016; Dahl et al., 2019; Plini et al., 2021). NA is released widely in cortical areas and participates in the modulation of sensory processing and many cognitive functions, such as attention, flexibility and memory (Sara, 2009; Sara and Bouret, 2012; Borodovitsyna et al., 2017; Poe et al., 2020; Breton-Provencher et al., 2021). The activation of the *Locus Coeruleus* is sensitive to novel and salient sensory stimuli (Berridge and Waterhouse, 2003). In the hippocampus, NA release can be induced by environmental enrichment (Galani et al., 2007; Brenes et al., 2009) and may contribute to spatial memory by enhancing long-term synaptic plasticity (Li et al., 2013). All these properties of the noradrenergic system suggest that soliciting this neuromodulatory system could be an effective strategy to promote better cognitive aging.

Our group previously reported that 30-40 days of olfactory enrichment (exposure to a new odor every day) is activating the noradrenergic system, inducing NA release and improving olfactory discrimination and memory in young adult mice (Veyrac et al., 2009; Rey et al., 2012). In contrast, it has no such effect on olfactory memory in aged mice (Rey et al., 2012). Based on this literature, we hypothesized that lifelong olfactory enrichment sessions would allow repeated activation of the noradrenergic system and improve cognition at late ages. An additional question that remains is to assess whether long-run olfactory enrichment would be efficient when started late in life. For that purpose, in the current study, we implemented olfactory enrichment sessions, starting in young adulthood (2 months) or at middle age (12 months) and that were repeated along the lifespan of the mice. We performed a longitudinal follow up, from young adulthood to 18 months, of olfactory performances, spatial memory and cognitive flexibility, as these are known to be age-sensitive cognitive functions (*e.g.* Roman et al., 1996; Barense, 2002; Schoenbaum et al., 2006; Murai et al., 2007; Brushfield et al., 2008; Wimmer et al., 2012). We then investigated the integrity of the *Locus Coeruleus* and the density of NA innervation in brain regions related to the assessed behavioral performances. Last, we characterized brain activation and functional connectivity through c-Fos mapping, to identify network remodeling associated to the cognitive performances. We report benefits of long-run olfactory enrichment in olfactory performances, spatial memory and cognitive flexibility, associated with a local increase of noradrenergic innervation and task-specific remodeling of neural networks.

## MATERIAL AND METHODS

### Animals

Adult male C57Bl/6J mice were housed in groups of 5-7 in standard laboratory cages and were kept on a 12 hours light/dark cycle, at a temperature of 22°C, with food and water ad libitum. All mice were purchased at the age of 2 months (Charles River laboratory, Domaine des Oncins, France) and aged in our animal facility. All efforts were made to minimize the number of animals used and their suffering during the experimental procedure in accordance with the European Community Council Directive 2010/63/EU. Protocols were approved by the French Ethical Committee (DR2013-48 (vM)).

### Experimental design

In order to assess the benefits of a long-run olfactory enrichment, two groups of mice were submitted to repeated sessions of olfactory enrichment: one starting at young adulthood (2 months, early-enriched group), the other one starting at middle age (12 months, late-enriched group). Each group of enriched mice (early enriched n=17; late enriched n=16) was compared to a control non-enriched group (early control n=15; late control n=16). The enrichment session lasted 10 days (5 weekdays during 2 consecutive weeks) and was repeated every month until mice reached 18 months (Fig. 1). Behavioral tests were performed to assess olfactory discrimination, spatial memory, associative learning and flexibility before, during and at the end of each enrichment period at 2, 12, 15 and 18 months for early groups, and 12, 15 and 18 months for late groups (Fig. 1). Behavioral testing took place during non-enriched periods. To investigate the effect of long-run olfactory enrichment on brain activation and connectivity, and the status of the noradrenergic system (*Locus Coeruleus* integrity and noradrenergic innervation), 18-month-old control and enriched mice were killed 1 hour after performing the olfactory discrimination task or the spatial memory task and processed for histological examination.

**Figure 1:**
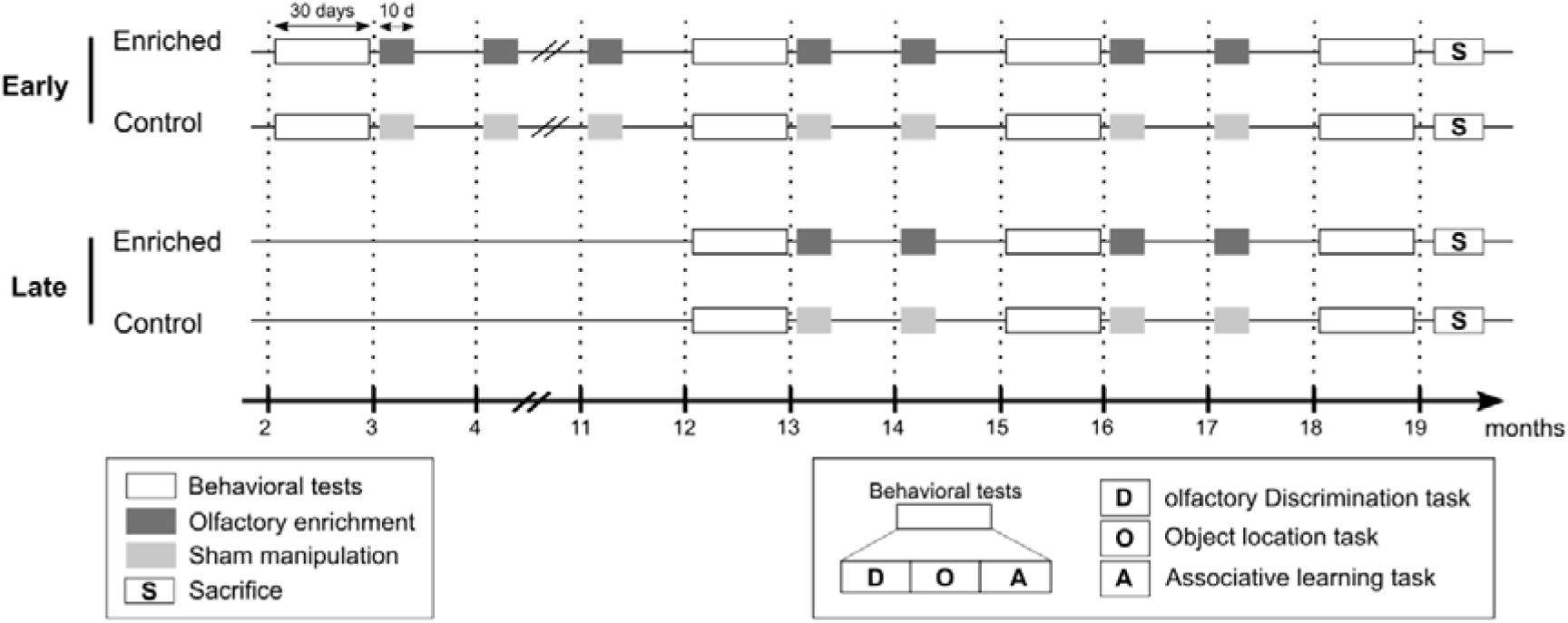
Experimental design. Mice were submitted to repeated sessions of olfactory enrichment starting at young adulthood (2 months, early group) or at middle age (12 months, late group). Each group of enriched mice was compared to a control non-enriched group (sham manipulation). Each enrichment session lasted 10 days (1h odor exposure/day, new odor everyday) and sessions were repeated every month until mice reached 18 months. Behavioral evaluation (olfactory discrimination, spatial memory, associative learning and cognitive flexibility) was performed before the beginning of the enrichment and at different time points during aging (12, 15 and 18 months). Mice were sacrificed (S) within 2-4 days after the last enrichment session. Subsets of 18-month mice performed the olfactory discrimination task or the object location task 1 hour before sacrifice.

### Olfactory enrichment

Enrichment sessions took place in the morning (between 8am and 1pm), in a ventilated cabinet and lasted 1h per day for 10 days. For the enriched group, an odor was placed in 2 tea balls hanging from the lid of the cage, as opposed to the control non-enriched group where the tea balls were empty (sham manipulation). Enrichment odors were 10 natural odors: white pepper, star anise, cinnamon, chocolate, cardamom, fennel, garlic, ginger, nutmeg and clove. A different odor was diffused each day. The same set of 10 odors was used for each monthly enrichment sessions. The order of odors’ presentation was randomized during the different enrichment sessions.

### Behavior

All behavioral evaluations were performed blind with regard to the enrichment status of the mice and conducted in the afternoon (12pm–7pm).

#### Olfactory discrimination

##### Behavioral testing

We used a cross-habituation task (Cleland et al., 2002) to assess spontaneous discrimination between a pair of perceptually similar odorants. This task was previously described (Mandairon et al., 2006, 2018; Vinera et al., 2015), including in aged mice (Rey et al., 2012; Moreno et al., 2014; Greco-Vuilloud et al., 2022). A session consisted of a habituation phase, during which the same odorant of the pair was presented 4 times (habituation trials: OHab1-OHab4, 50-second presentations with a 5-minute intertrial interval), followed by a discrimination phase, during which a new odorant (the other odorant of the pair) was presented (test trial: OTest, 50-second presentation) (Fig. 2A). Each mouse underwent two sessions, during which each odorant of the pair was alternatively used as habituation or test odorant. Before the habituation phase, a non-odorized tea ball (containing odorless mineral oil) was presented to the animal. The odorant was presented in a tea ball (60µl odorant at a vapor phase pressure of 1Pa, diluted in mineral oil and spotted on a filter paper), hanging from the cover of the cage. The amount of time that the mice spent investigating the odorant was recorded manually. Investigation was defined as active sniffing within 1cm of the tea ball. Habituation resulted in a decrease in the investigation time across the four successive presentations of the same odorant (OHab1 to OHab4). During the last trial (OTest), an increase of investigation time indicated discrimination between the two odorants (OTest > OHab4), while similar or shorter investigation time indicated that the mouse did not discriminate between the two odorants (OTest ≤ OHab4).

**Figure 2:**
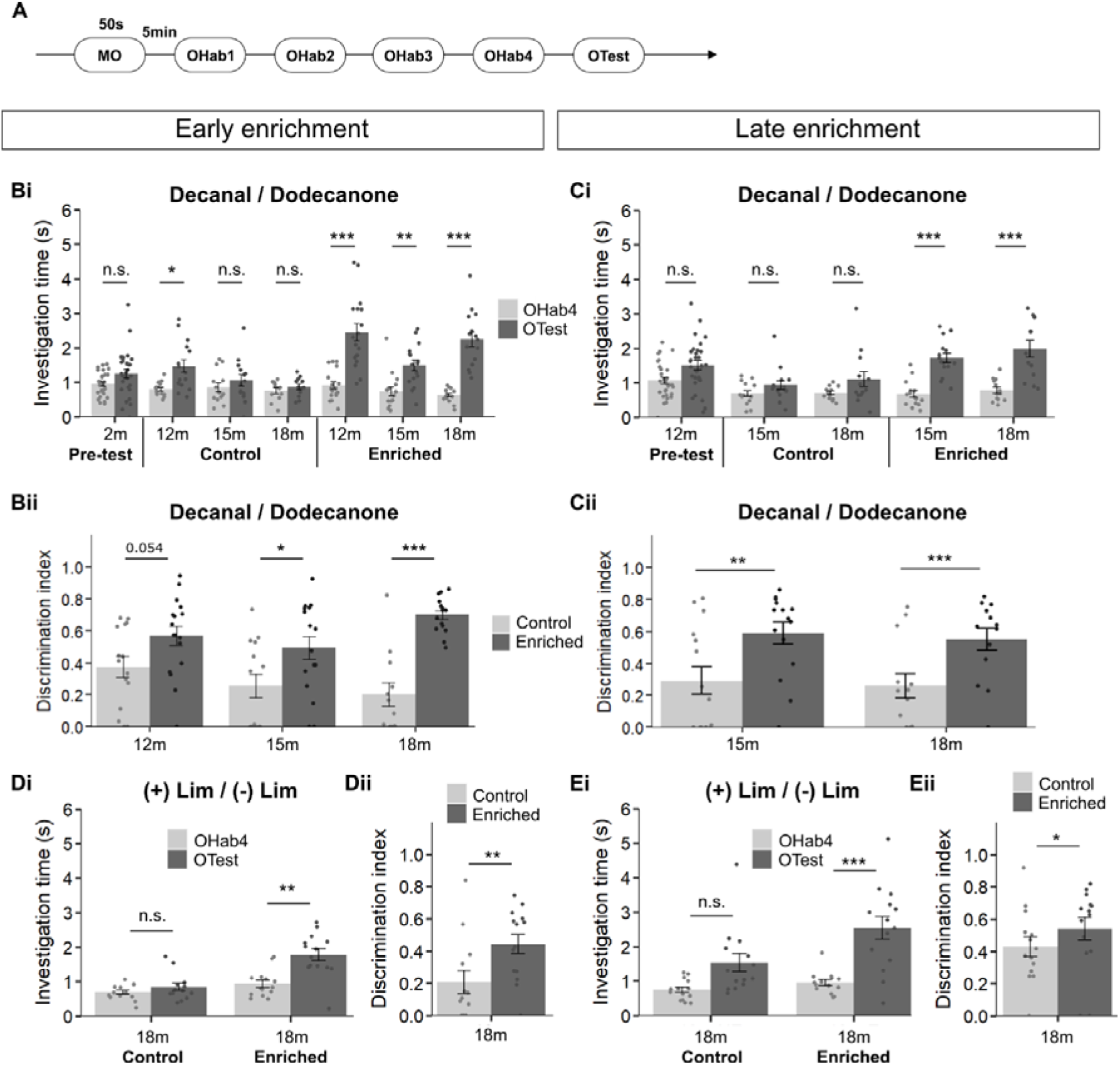
Fine discrimination abilities are improved by long-run olfactory enrichment until late ages. **(A)** Assessment of fine olfactory discrimination at different ages in the enriched and control groups of the early- and late-started enrichment, using a cross habituation task. **(B-C)** Discrimination between Decanal / Dodecanone or **(D-E)** (+) Limonene / (-) Limonene. Data are presented as mean investigation time (Bi, Ci, Di and Ei) or mean discrimination indices (Bii, Cii, Dii and Eii) ± standard error of the mean. Dots represent data from individual mice. n= 12-28 mice per group (see Table 2 for details). * p<0.05; ** p<0.01; *** p<0.001. MO =mineral oil.

##### Odorants

4 odorants were used: Decanal (CAS #112-31-2, purity >98%, Sigma Aldrich), 2-Dodecanone (CAS #6175-49-1, purity >97%, Aldrich Sigma), (+) Limonene (CAS #5989-27-5, purity >97%, Sigma Aldrich) and (-) Limonene (CAS #5989-54-8, purity >97%, Sigma Aldrich). Based on our previous studies (Moreno et al., 2009; Rey et al., 2012), these odorants were selected to compose two pairs of perceptually similar odorants that are not spontaneously discriminated by mice (Decanal/Dodecanone, (+)Limonene/(-)Limonene). The mice were tested for discrimination with each pair on separate days and the order of testing of these pairs was randomized during the behavioral sessions.

##### Data analysis

For each mouse and trial, investigation time was averaged between the two sessions of the same pair of odorants (each odorant of the pair was alternatively used as habituation or test odorant). Then, investigation time was averaged between mice within groups for each trial. Individual sessions with investigation times lower than 0.5s at OHab1 were excluded, as well as sessions where one trial exceeded mean +/-2SD (these exclusions represented 187 sessions among a total of 957 sessions). Removing the whole session rather than the single trial allowed performing paired statistical analysis. We calculated a discrimination index defined as [1-(OHab4/OTest)], where OHab4 and OTest were the exploration time during respectively OHab4 and OTest. The discrimination index approaches 0 when OTest is similar to OHab4 (no discrimination) and 1 when OTest largely exceeds OHab4. When OHab4 was superior to OTest, the index was set to 0. This index enables the comparison of discrimination performances independently of overall variations in investigation time and provides a graded measure of discrimination while the cross-habituation task is a binary evaluation (Cleland et al., 2002; Rey et al., 2012). To compare investigation times, as the data did not satisfy to normality and variance equality criteria (assessed with Shapiro-Wilk normality test on residues and Bartlett test of homogeneity of variances), we performed non-parametric tests: a Friedman’s ANOVA to test for trial effect (OHab1-4, OTest) followed by pairwise Nemenyi tests corrected for multiple comparisons (OHab1 vs. OHab4 to test for odor habituation and OHab4 vs. OTest to test for odor discrimination). To compare indices between control and enriched mice, 2-way ANOVA (type III ANOVAs with enrichment and age as factors) were performed and followed by unilateral pairwise t.tests with Holm correction for multiple comparisons when data respected normality and variance equality criteria, otherwise unpaired unilateral two samples Wilcoxon tests were performed. When indices data respected normality but not variance equality, unilateral pairwise t.tests with Welch correction and Holm correction for multiple comparisons were used. Detailed description of the statistical results is provided in Table 2. Data were analyzed using R-Studio® (stats package and PMCMRplus package for Friedman’s ANOVA). The level of significance was set to 0.05.

#### Associative and flexibility capacities

##### Behavioral testing

We used an olfactory associative learning task (Mandairon et al., 2009, 2018) with rule reversal (Fig. 3A), as described previously (*e.g.* Guérin et al., 2009; Sultan et al., 2010). Task reversal is a widely used paradigm to evaluate cognitive flexibility in animal models (Izquierdo et al., 2017).

**Figure 3:**
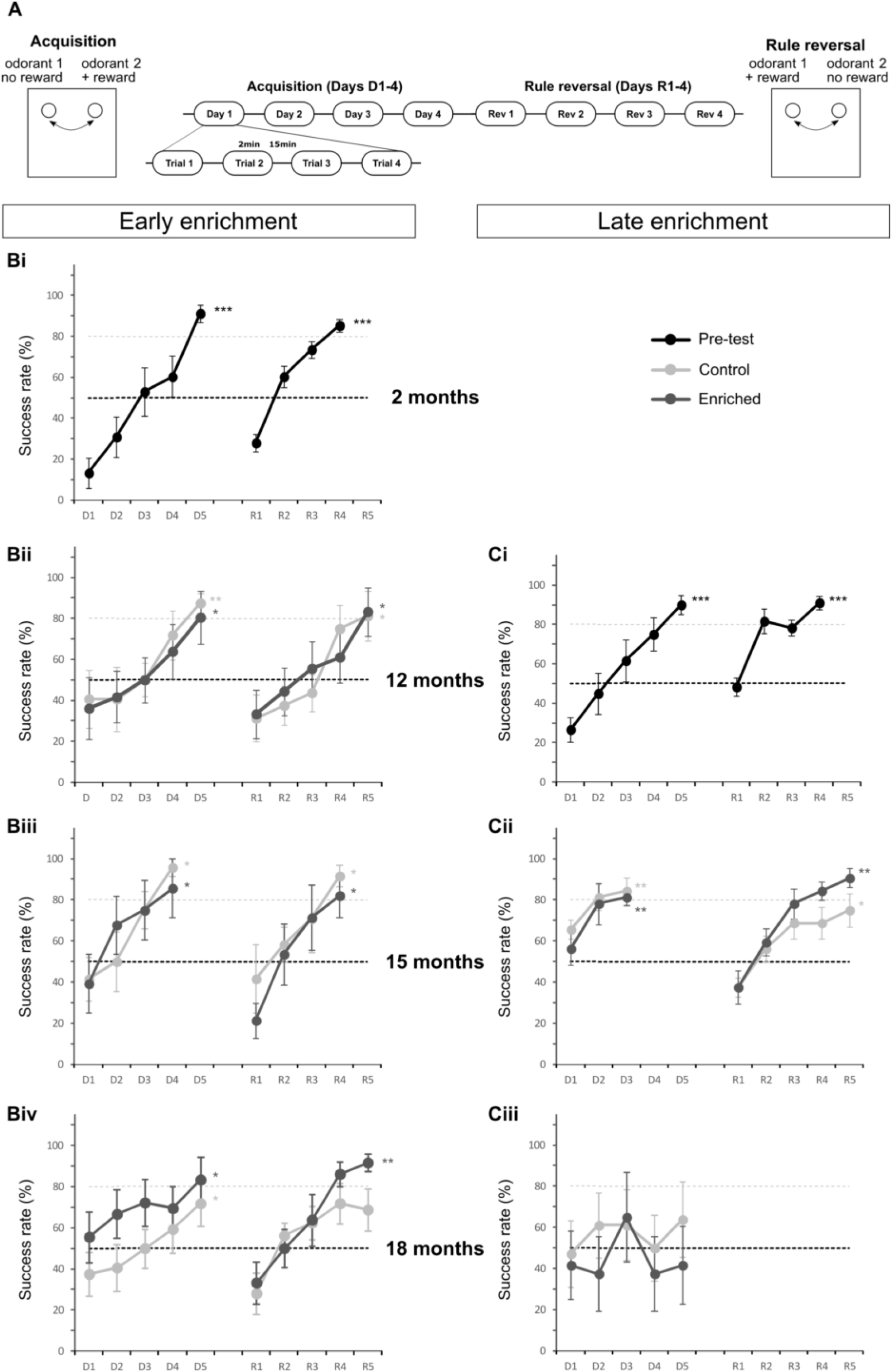
Associative learning and cognitive flexibility upon long-run olfactory enrichment. **(A)** Associative learning and cognitive flexibility were evaluated using an olfactory associative task during the acquisition of the first rule (D1-D5) and after the reversal of the rule (R1-R5). **(B)** Mice of the early group were tested at 2 months before the beginning of the enrichment (pre-test) and at different ages (12, 15 and 18 months) in the enriched and control groups. **(C)** Mice of the late group were tested at 12 months before enrichment (pre-test) and then during enrichment at 15 and 18 months. Data are presented as mean success rate ± standard error of the mean. n= 6-17 mice per group (details are provided in Table 3). * p<0.05; ** p<0.01; *** p<0.001 for comparisons to 50% (chance level).

##### Shaping

One day before the beginning of the experiment, the mice were food deprived (2g per day, leading to a maximum of 20% reduction in body weight). Shaping consisted in sessions of 4 trials (trial duration = 2min, one session per day) during which mice were trained to retrieve a reward (a small bit of cereal, Corn Flakes, Kellogg’s®) hidden in one hole of a two-holeboard apparatus, in the absence of any odorant. During the first trials, the reward was placed on top of the bedding. After the mice retrieved the reward, it was buried deeper and deeper in the bedding. The shaping was considered as completed when every mouse successively retrieved the buried reward (2-3 sessions) in 80% of the trials, and was followed by the conditioning phase.

##### Conditioning

Conditioning consisted in sessions of 4 trials (trial duration = 2min, intertrial interval = 15min, one session per day), during which the two holes were odorized with perceptually dissimilar odorants (see below) During acquisition, one of the odorants was systematically reinforced and mice had to retrieve the reward. To avoid spatial learning, the location of the odorants was randomized between the two holes. A control pseudo-conditioned group was also constituted, in which the reward was randomly placed in either of the odorized holes, so that the mice would not associate a specific odorant with the reward. For each trial, correct choices (first nose poke in the reinforced hole) were recorded as indicative of learning. We calculated a percentage of correct choices for each session. The learning criterion was set at 80% of correct choices for the group. The rule was reversed once the group had reached this criterion: the previously non-reinforced odorant became the reinforced odorant. The same protocol and the same parameters were used for the conditioning with the modified rule.

##### Odorants

We used three pairs of dissimilar odorants: (+) Limonene (CAS #5989-27-5) and Decanal (CAS #112-31-2), Isoamyl Acetate (CAS #123-92-2, purity >95%, Sigma Aldrich) and Anisole (CAS #100-66-3, purity >98%, Sigma Aldrich), 3-Hexanol (CAS #623-37-0, purity >97%, Fluka, Buchs, Switzerland) and Heptanal (CAS #111-71-7, purity >95%, SAFC, Sigma Aldrich) (Table 1). 20µl of pure odorant were spotted on a swab, that was deposited in the hole and covered with bedding. We used three pairs of odorants in order to distribute odorants between testing at different ages and thereby minimize repetitions. In order to limit the risk of over-training, mice that were pseudo-conditioned during one session (ex. at 12 months) were conditioned on the next session (ex. at 15 months) and vice-versa. At each age, control and enriched mice were tested with the same pair of odorants and the same odorant was reinforced in the conditioning phase before reversal.

**Table 1:** Odorants used in the olfactory associative learning task.

|  | EARLY ENRICHMENT | LATE ENRICHMENT |
| --- | --- | --- |
| **2 months** | Amyl Acetate/ Anisol ;<br>1<sup>st</sup> rule : Amyl Acetate reinforced,<br>reversal : Anisol reinforced | — |
| **12 months** | 3 odor pairs ;<br>odor pairs and reinforced odor<br>were randomized within groups | Amyl Acetate / Anisol ;<br>1<sup>st</sup> rule : Amyl Acetate reinforced,<br>reversal : Anisol reinforced |
| **15 months** | 3 odor pairs ;<br>odor pairs and reinforced odor<br>were randomized within groups | + Limonene / Decanal ;<br>1<sup>st</sup> rule : + Limonene reinforced,<br>reversal : Decanal reinforced |
| **18 months** | 3 odor pairs ;<br>odor pairs and reinforced odor<br>were randomized within groups | 3-Hexanol / Heptanal ;<br>reinforced odor was<br>counterbalanced within groups |

##### Data analysis

For each behavioral session, we calculated a mean percentage of correct choices for each group. Shapiro and Fisher tests were performed to assess data normality and variance homogeneity for each group and each session. As these conditions were not respected, we performed non-parametric statistical analysis. We used Friedman followed by Nemenyi *post hoc* tests (R software, PMCMRplus package) in order to compare the percentage of correct choices across days. Univariate Wilcoxon tests (R software, stats package) were used to evaluate whether the success rate was higher than 50% (chance level). Detailed description of the statistical results is provided in Table 3.

#### Spatial memory

##### Behavioral testing

To assess spatial memory, we used the object location test inspired by previous works (*e.g.* (Murai et al., 2007; Wimmer et al., 2012; Cès et al., 2018). This task is based on the spontaneous tendency of rodents to preferentially re-explore a displaced object when it is confronted to a non-displaced object and recognizes that it has been displaced. The test was conducted in an open field (length 50x width 50x height 30cm) containing a pair of objects (objects A and B). Three different pairs of objects were used: two black metal towers, two Lego® blocks and two Playmobile®, all approximately of the same volume (5x5x2cm) but pairs differed in shape and colors. A visual cue was present on one of the four walls of the experimental arena. After familiarization to the experimental chamber (5min per day during 2 consecutive days in the empty arena), the task consisted in an acquisition trial and a test trial (Fig. 4A). The acquisition trial consisted in free exploration of the objects in the arena during 5min. Each object was placed in opposite corners of the open field (at a distance of 12.5cm from the walls). At the end of the acquisition trial, the mice were returned to their home cage. After a delay of 5, 15 or 60min, the mice were reintroduced in the arena for the retention trial (duration 5min) after one object had been moved to another corner of the apparatus (displaced object, DO) while the other object remained at the same location (non-displaced object, NDO). The displacement distance was 24cm. Each mouse was tested for all three delays with different pairs of objects and on separate days.

**Figure 4:**
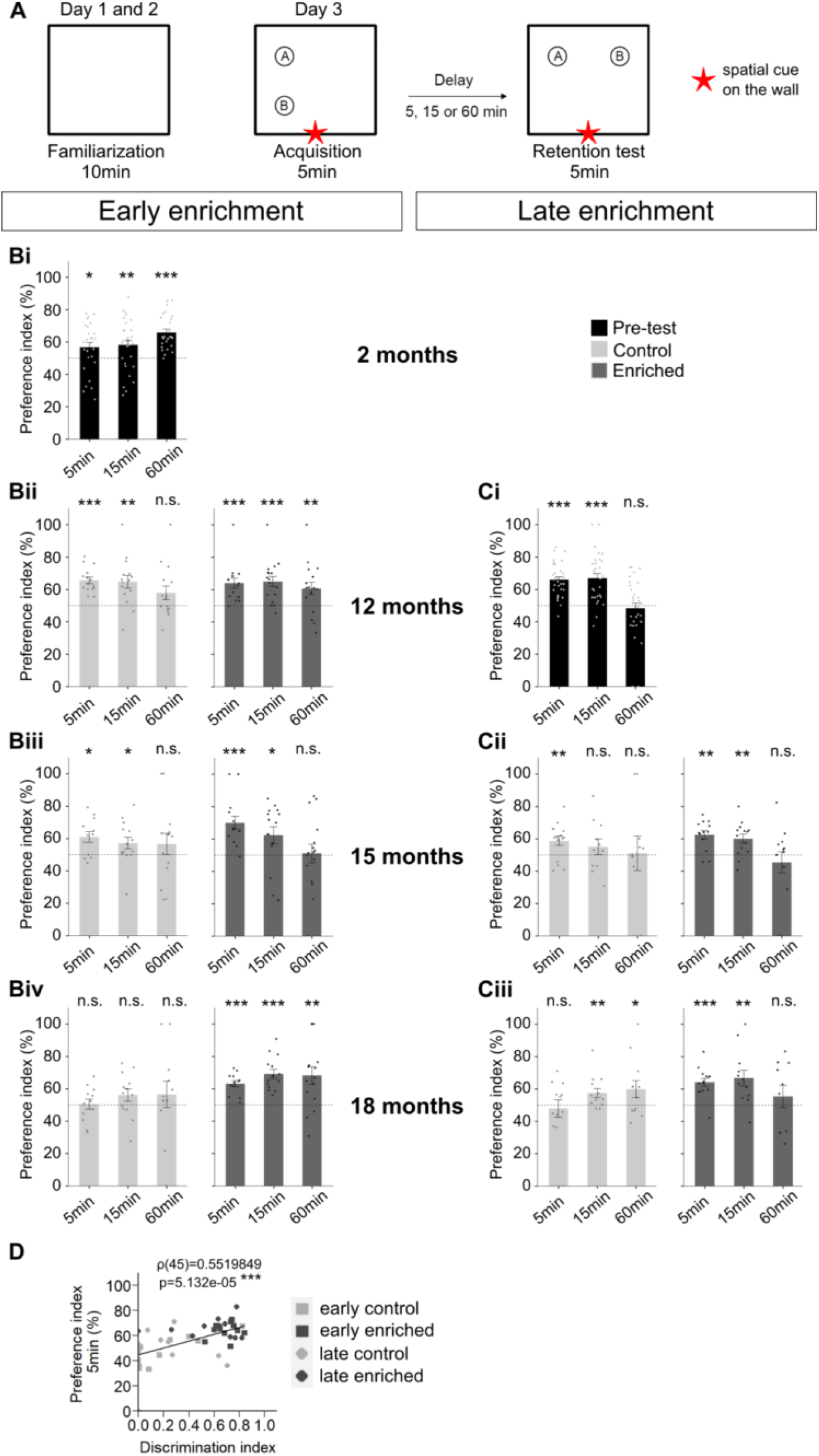
Spatial memory is improved by long-run olfactory enrichment. **(A)** Schematic of the procedure used in the object location task. **(B)** Preference index for pre-test at 2 months and during enrichment at 12, 15 and 18 months for the early enriched and control groups. **(C)** Preference index for pre-test at 12 months and during enrichment at 15 and 18 months for late enriched and control groups. A preference index significantly greater than 50% indicates that mice remember the initial position of the object. Data are presented as mean preference index ± standard error of the mean. Dots represent data from individual mice. n=12-32 mice per group (details are reported in Table 4). **(D)** Significant positive linear correlation was found between the discrimination index for the Decanal / Dodecanone pair and the preference index for the 5-min-delay at 18 months. n= 45 mice. * p<0.05; ** p<0.01; *** p<0.001.

Object location, new location, pairs of objects used and retention delays were counterbalanced over groups and sessions. During both acquisition and retention trials, we measured the investigation time of the two objects and calculated a preference index for each mouse. This index corresponds to the ratio of the investigation time of the displaced object to the total time spent exploring both objects (*i.e.* preference = 100*DO/(NDO+DO). We verified that the two objects were explored at similar levels during acquisition and we present here preference index for the retention trials.

##### Data analysis

A mean preference index was calculated for each group of mice and each delay. Shapiro and Fisher tests were performed to assess data normality and variance homogeneity. As these conditions were not respected, we performed non-parametric statistical analysis. Univariate Wilcoxon tests (R software, stats package) were used to compare the preference index to 50% (= no preference corresponding to similar exploration of the two objects). Detailed description of the statistical results is provided in Table 4.

### Histological evaluations

Histological evaluations were conducted on a subset of randomly-selected 18-month old mice and blind with regard to the enrichment status of the mice.

#### Tissue preparation

Mice were deeply anesthetized (pentobarbital, 0.4g/kg) and killed by intracardiac perfusion of paraformaldehyde (4% in phosphate buffer, pH 7.4). Brains were removed, postfixed, cryoprotected in sucrose (20%), frozen rapidly, and then stored at -20°C. Serial coronal sections (14μm thick) were obtained at -20°C on a cryostat HM 550 Microm (Microtech, Francheville, France).

#### *Locus Coeruleus* integrity

##### Phox2a/Tyrosine Hydroxylase (TH) immunohistochemistry

Sections were rehydrated in 1X Phosphate Buffer Saline (PBS), then permeabilized for 30min in Triton X-100 (0.5% in 1X PBS) and rinsed 5min in 1X PBS at room temperature (RT). To block nonspecific binding, sections were incubated at RT during 120min in a blocking solution containing horse serum (5%), goat serum (5%), bovine serum albumin (2%), Triton X-100 (0.1%) in 1X PBS. They were then incubated with mouse anti-phox2a (1:500, ref WH0000401M1-100UG, Merck) and a chicken anti-TH (1:1000, ref ab76442, Inst.J.Boy) antibodies diluted in the blocking solution during 48h at 4°C. After rinsing, sections were incubated for 120min at room temperature with biotinylated horse anti-mouse (1:200, ref A-2001, Vector) and goat anti-chicken Alexa 633 (1:250, ref A21103, Molecular Probes) secondary antibodies in a solution containing horse serum (5%) and goat serum (5%) in 1X PBS. Sections were rinsed and incubated for 90min with Streptavidin Alexa 488 (1:1000, ref S32354, Molecular Probes). Finally, after rinsing, sections were coverslipped with a Vectashield medium containing DAPI (Vector Lab).

##### Phox2a/TH-positive cells quantification

Double-labeled sections were examined using a pseudo-confocal microscope equipped with an ApoTome system, coupled with an Axio Vision 4.8 image analysis software (Zeiss microscope Axio Imager.ZI). z-stack images were acquired with a x20 objective (z-step = 2μm. *Locus Coeruleus* was delineated on each section with ImageJ software to determine its surface, based on Phox2A staining. Phox2a and TH-positive cells were manually counted in the anterior part of the *Locus Coeruleus* (2-6 sections/animal, distance from Bregma: -5.2 to +5.51mm), containing most of the neurons projecting to the anterior brain (Sara and Bouret, 2012; Chandler et al., 2014; Schwarz and Luo, 2015; Schwarz et al., 2015; Kebschull et al., 2016; Poe et al., 2020; Breton-Provencher et al., 2021). The number of Phox2a and TH-positive cells was divided by the surface of the *Locus Coeruleus* to yield the total density of labeled cells (labeled profiles/mm2) and an averaged density was calculated for each animal and then for the group (n=10-11 animals/pooled group). We also assessed the ratio of Phox2a neurons expressing the TH phenotype by yielding the percentage of TH-positive neurons over the whole Phox2a-positive population (n=10-11 animals/pooled group).

##### TH intensity

The mean gray value of TH immunostaining was assessed with ImageJ on delineated *Locus Coeruleus* areas. An averaged mean gray value was calculated for each animal (2-6 sections/animal) and then for the group (n=10-11 animals/pooled group).

##### Data analyses

All data were analyzed using R-Studio® (stats package and car package for ANOVAs). The level of significance was set to 0.05. As data showed normal distribution and variance equality (assessed on Shapiro-Wilk normality test on residues and Bartlett test of homogeneity of variances), statistical analysis was carried out by 2-way ANOVA (type III ANOVA) with enrichment and onset of the enrichment (early or late) as factors (detailed statistical results are provided in Table 5). ANOVAs did not show significant effect of the onset of the enrichment nor interaction with enrichment and early and late controls were pooled, as well as early and late enriched groups. Then, pooled controls and pooled enriched mice were compared by unilateral Student t.test or unilateral unpaired Wilcoxon test. Detailed description of the statistical results is provided in Table 5.

#### Noradrenergic innervation

##### NET (NorEpinephrin Transporter) immunohistochemistry

Sections were rehydrated in 1X Phosphate Buffer Saline (PBS), then permeabilized for 30min in Triton X-100 (0.5% in 1X PBS) and rinsed 5min in 1X PBS at room temperature. To block nonspecific binding, sections were incubated at RT during 90min in a blocking solution containing horse serum (5%), bovine serum albumin (2%), Triton X-100 (0.1%) in 1X PBS. They were then incubated with mouse anti-NET primary antibody (1:1000, ref AMAB91116-100UL, Sigma Aldrich) diluted in the blocking solution during 72h at 4°C. After rinsing, sections were incubated for 120min at RT with biotinylated horse anti-mouse (1:200, ref A-2001, Vector) secondary antibody in a solution containing horse serum (5%) in 1X PBS. Sections were rinsed and incubated for 90min with Streptavidin Alexa 488 (1:1000, ref S32354, Molecular Probes). Finally, after rinsing, sections were coverslipped with a Vectashield medium containing DAPI (Vector Lab).

##### Image acquisition and quantification of NA innervation

3D images were acquired with a pseudo-confocal microscope equipped with an ApoTome system, coupled with an Axio Vision 4.8 image analysis software (Zeiss microscope Axio Imager.ZI). To quantify the area occupied by the NA fibers, z-stack images were acquired with a x40 immersion objective (z-step = 1μm). Pictures were randomly taken on 2 different sections for each animal within the granule cell layer to sample the OB, the stratum oriens, the pyramidal layer and the stratum radiatum to sample cornu ammonis CA1, and the molecular and granule cell layers to sample the dentate gyrus (DG). The stained surface was measured on a maximum intensity projection of the z-stack. The stained surface was automatically measured by applying the same intensity threshold to all images with ImageJ software. Results are expressed as the ratio between the NET-labeled area and the total area of the image (% of stained surface). An average stained surface is calculated for each animal and then for the group (n=8-12 animals/group).

##### Data analyses

All data were analyzed using R-Studio® (stats package and car package for ANOVAs). The level of significance was set to 0.05. When data showed normal distribution and variance equality (assessed on Shapiro-Wilk normality test on residues and Bartlett test of homogeneity of variances), statistical analysis was carried out by 2-way ANOVA (type III ANOVA) with enrichment and onset of the enrichment (early or late) as factors (detailed statistical results are provided in Table 6). When normality was confirmed but not variance homogeneity, bilateral Student t.tests with Welch correction and with Holm correction for multiple comparisons were performed. When both normality and variance homogeneity were not confirmed, bilateral unpaired Wilcoxon tests with Holm correction for multiple comparisons were performed. None of these analyses revealed significant difference between early and late groups and these were pooled and comparisons were made between pooled controls and pooled enriched groups with unilateral unpaired Student t.test or unilateral unpaired Wilcoxon test. Detailed description of the statistical results is provided in Table 6. For linear correlations, as data did not respect normality, we performed unilateral Spearman tests.

#### Brain activation pattern and connectivity analysis

To investigate the neuronal networks activated during a specific task, subgroups of control and enriched mice were constituted based on the behavioral task they performed 1h before the sacrifice: one performed the olfactory discrimination task using a pair of perceptually similar odorants (Decanal and 2-Dodecanone (early- and late-enriched and control mice) and the other the object location task with a 60-min delay (early-enriched and control mice). This procedure allows to identify neurons activated during the task by the expression of the immediate early-gene c-Fos (Guzowski et al., 2005). Network analyses for the discrimination task sub-group were performed by pooling the early and late-enriched groups (n=7), as well as their respective control groups (n=7). For the object location task sub-group, analyses were performed on the early-enriched and control groups (n=5 and n=5 respectively).

##### c-Fos immunohistochemistry

Sections were rehydrated in 1X Phosphate Buffer Saline (PBS), then permeabilized for 30min in Triton X-100 (0.1% in 1X PBS) and rinsed in 1X PBS at RT. To block endogenous peroxidases, sections were incubated in H2O2 (3%) and rinsed in 1X PBS. To block nonspecific binding, sections were incubated at RT during 90min in a blocking solution containing goat serum (5%), bovine serum albumin (2%), Triton X-100 (0.5%) in 1X PBS. They were then incubated with rabbit anti-c-Fos (1:2000, ref 26 192-AP, ProteinTech) primary antibody diluted in the blocking solution during 48h at 4°C. After rinsing, sections were incubated for 120min at RT with biotinylated goat anti-rabbit secondary antibody (1:200, ref BA-1000, Vector) in a solution containing goat serum (5%) and BSA (5%) in 1X PBS. Sections were rinsed and incubated for 60min with an avidine-biotine peroxidase kit (Vectastain Elite kit, Vector Laboratories, Burlingame, CA, USA). After rinsing, sections were pre-incubated in a diaminobenzidine solution (0,06% in Tris HCl 0,05M) and revealed with the diaminobenzidine solution containing NiCl2 (0,03%) and H2O2 (0,06%).

##### c-Fos-positive cell quantification

For each section analyzed, the external contour of the right hemisphere (or the left when the right was occasionally unavailable) was outlined and c-Fos-positive cells were automatically counted using a mapping software (Mercator, Explora Nova, La Rochelle, France) coupled to a Zeiss microscope. The coordinates of each labelled cell and the contour point were exported into a software made by the team allowing high-accuracy matching of experimental brain sections with a reference brain atlas (Midroit et al., 2018). This method allows precise, automatic assignment of the labelled cells to a brain structure and permits between-group comparisons after re-fitting in a common anatomical space. A total of 31 areas was analyzed. These areas were distributed into 6 functional systems named olfaction, prefrontal, spatial memory, motor, other sensory and neuromodulatory. Ten areas were selected based on their known involvement in olfactory processing and constituted the “olfaction” system: the glomerular (GL) and granule cell (GrL) layers of the OB, the accessory olfactory bulb (AOB), the anterior olfactory nucleus (AON), tenia tecta (TT), piriform cortex (Pir), tubercle (Tu), cortical amygdala (ACo), nucleus of the lateral olfactory tract (LOT), postero-lateral cortical amygdala (PLCo). The “memory” system (7 areas) was composed of lateral (LSN) and medial (MSN) septal nucleus, entorhinal cortex (EC), dorsal dentate gyrus (dDG), dorsal CA3 (dCA3), dorsal CA1 (dCA1) and lateral habenula (LHb). The “prefrontal” system (7) was composed of the cingular cortex (CC), frontal association cortex (Fr), insular cortex (IC), lateral (LO), medial (MO), ventral (VO) and dorso-lateral (DLO) orbital cortex. Last, 2 areas composed the “motor system” (motor cortex (MC), caudate and putamen (CPu)), 3 areas composed the “other sensory systems” (parietal cortex (PC), somatosensory cortex (S1), auditory cortex (AC)) and 2 composed the “neuromodulatory system” (diagonal band of Broca (HDB) and *Locus Coeruleus* (LC)). Missing sections were interpolated, as explained in Midroit et al., 2018).

##### Comparison of cell densities

In order to reduce inter-individual variability in staining intensity, c-Fos density for each area of a given animal was normalized by the average c-Fos density of all 31 regions of this animal. Results from enriched animals were then expressed as a percentage of change relative to controls.

##### Functional connectivity analysis and network construction

Functional connectivity analysis was performed similarly to previous studies using the graph theory (Wheeler et al., 2013; Babayan et al., 2017; Vetere et al., 2017). Within each experimental group (olfactory discrimination task: controls and enriched, object location task: controls and enriched), pairwise correlations between non-normalized c-Fos densities in the 31 regions were determined by computing Pearson’s correlations (totaling 465 pairs of regions). Each set of correlations was displayed as color-coded correlation matrices using R software. Networks graphs were constructed by considering only correlations reaching a bilateral significance level of p ≤ 0.025 corresponding to r ≥ 0.82 (uncorrected for multiple comparisons). We verified that the main observations were not dependent on the threshold and were retrieved with other thresholds of p ≤ 0.05, p ≤ 0.01 and p≤ 0.005. Each region is represented by a point called a node. The lines between the regions represent the correlations. These correlations represent the co-variation of the density of c-Fos-positive cells in two regions across mice and are interpreted as functional connections between these two regions (Horwitz et al., 1995; Wheeler et al., 2013; Babayan et al., 2017; Vetere et al., 2017). We also calculated the connection density defined as the proportion of significant correlations (=effective connections) in the whole network or in a functional system to all possible connections. Finally, we assess the load of each system among effective connections as the proportion of connections of one functional system to all effective connections.

##### Measure of networks parameters

Complex networks can be both segregated – enabling for specialized processing in more densely connected clusters – and integrated – enabling for efficient information flow through the network – (Bullmore and Sporns, 2009). Networks measures were generated using functions from Olaf Sporns’ brain connectivity toolbox (https://www.brain-connectivity-toolbox.net/). To assess the integration of our networks, we measured their global efficiency, defined as the average inverse shortest path length in the network (the shortest path length being the minimum number of nodes that must be traversed to go from one node to another) (Bullmore and Sporns, 2009). To assess the segregation, we computed the mean clustering coefficient that represents the average clustering coefficient of all nodes of the network. The clustering coefficient represents the number of connections that exist between the nearest neighbors of a node divided by the total number of possible connections between them (Rubinov and Sporns, 2010). Experimental network’s parameters were compared to those calculated from 1000 random networks, each random network sharing the same number of active nodes, connections and same degree distribution with the experimental network (Rubinov and Sporns, 2010). For each experimental network, we generated 95% confidence intervals by bootstrapping. Bootstrapping involves resampling subjects with replacement 1000 times and recalculating network measures as described in Wheeler et al., 2013. When the confidence intervals do not overlap, the parameters of the experimental network are considered different from the random network.

##### Cluster analysis

The cluster analysis was conducted with the Markov Cluster Algorithm (inflation parameter set at 2), a scalable, unsupervised cluster algorithm for networks based on simulation of stochastic flow in graphs (https://www.micans.org/mcl/). Clusters and associated visualizations were performed using Cytoscape (http://www.cytoscape.org/) (Wheeler et al., 2013; Babayan et al., 2017).

##### Data analyses

c-Fos densities and connection densities were analyzed using in-house made scripts written in R-Studio® (stats package and car package for ANOVAs). The level of significance was set to 0.05. Shapiro-Wilk and Bartlett preliminary tests were performed to assess respectively the normality and homoscedasticity of the data. For comparison to a mean of reference (100%), we performed bilateral one-sample t.tests corrected for multiple comparisons (Fig. 7 and Table 7). For proportion comparisons, we performed bilateral 2-sample tests for equality of proportions (Fig. 8).

## RESULTS

In order to assess the sensory and cognitive benefits of a long-run olfactory enrichment, mice were submitted to repeated sessions of olfactory enrichment starting at young adulthood (2 months) and compared to a control non-enriched group (Fig. 1). In addition, to evaluate if a late-started enrichment could also be beneficial, another group was submitted to enrichment starting at middle age (12 months) and compared to its respective control non-enriched group. Olfactory discrimination, olfactory associative memory, cognitive flexibility and spatial memory were assessed at 2, 12, 15 and 18 months for the early groups and at 12, 15 and 18 months for the late groups.

### Long-run olfactory enrichment improves olfactory discrimination at late ages, whether started early or late in life

Olfactory discrimination performances were evaluated using a cross-habituation task with a pair of perceptually similar odorants (Decanal and 2-Dodecanone) (Fig. 2A). Significant habituation to the odorants was observed in all groups, as revealed by the decrease in investigation time from OHab1 to OHab4 (Supp. Fig. S1 and Table 2 for complete statistical report). Discrimination results are depicted in Fig. 2, by comparing the investigation time of the test odorant (OTest) to the fourth presentation of the habituation odorant (OHab4).

**Table 2:**
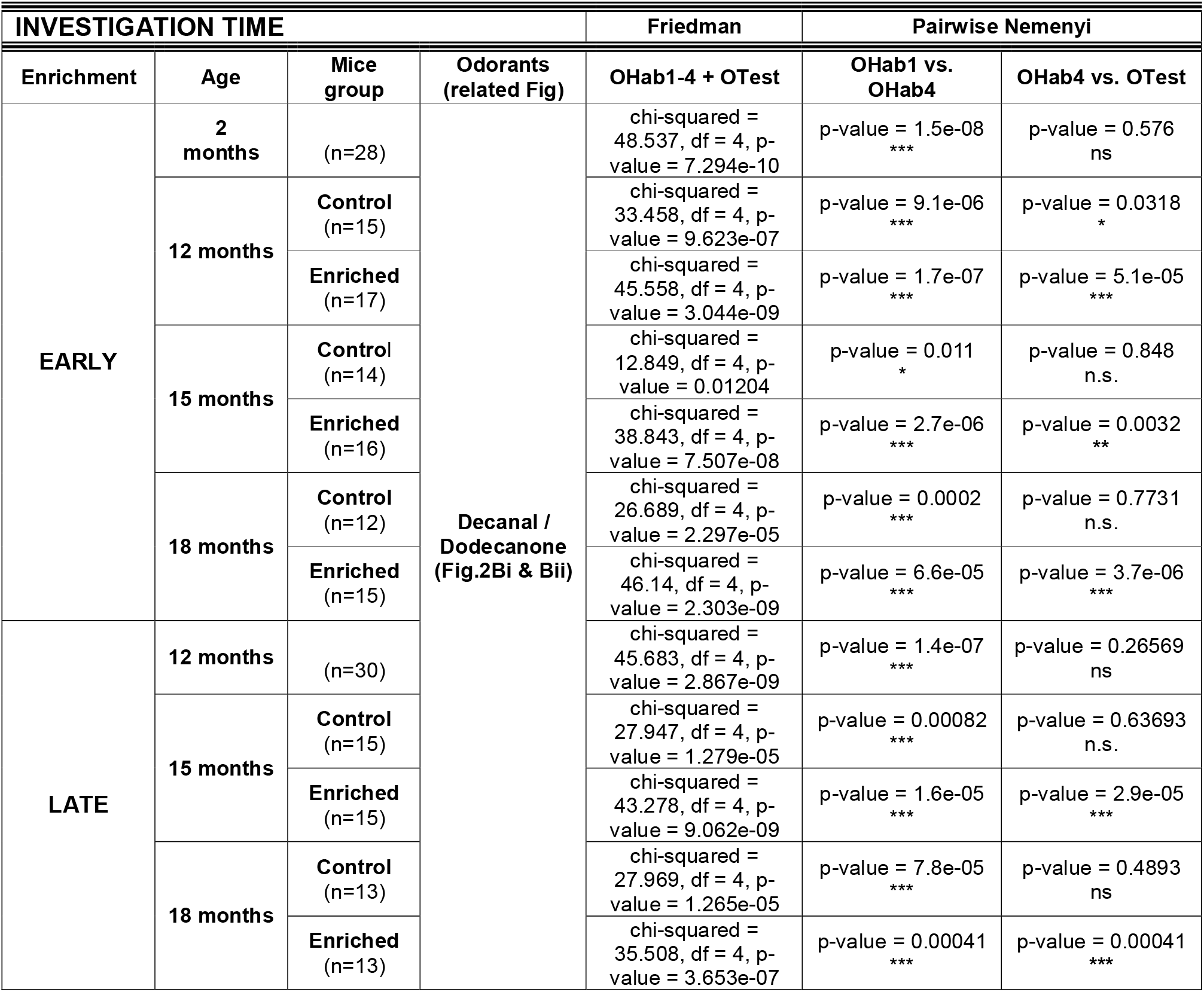
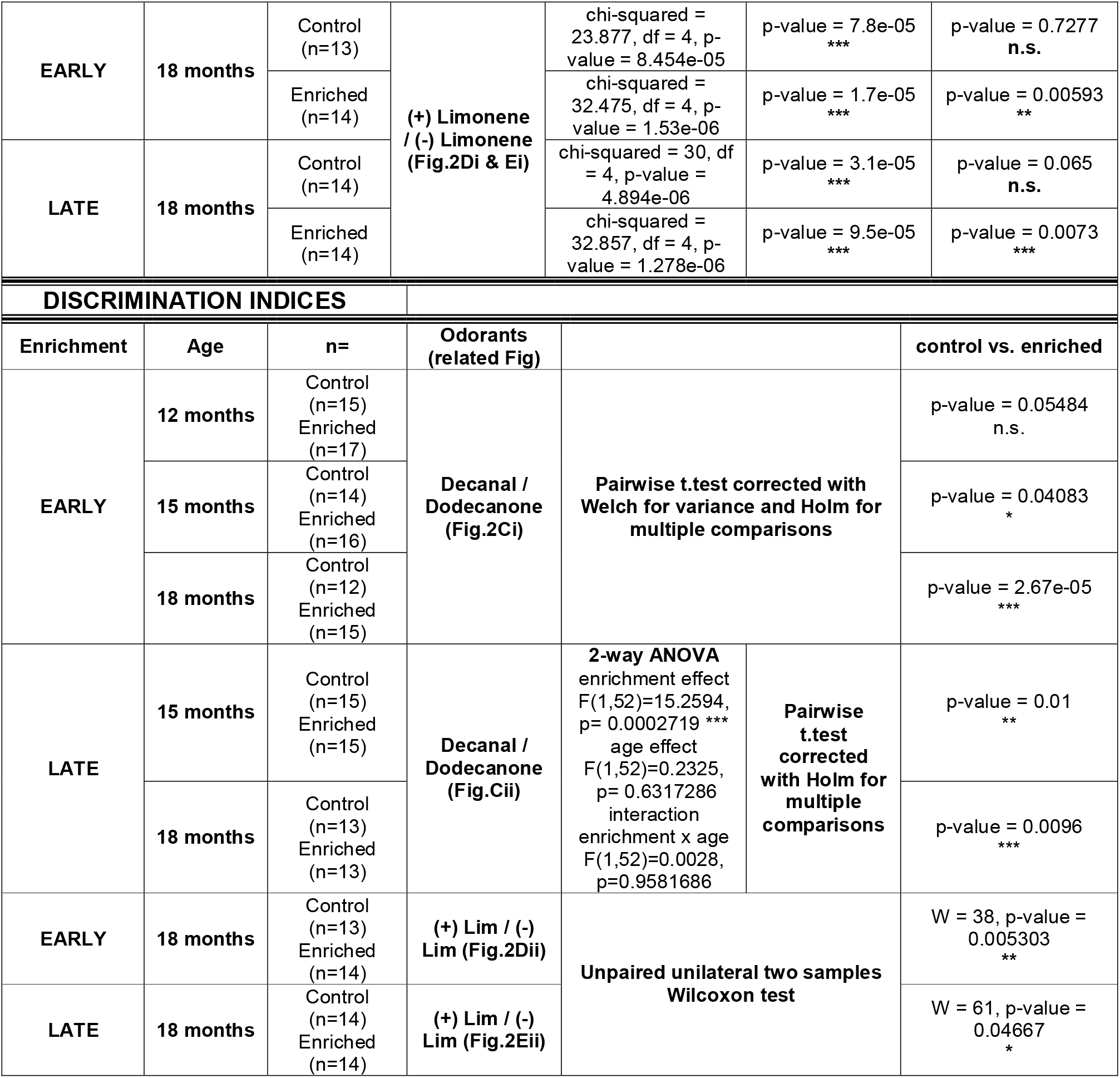
Detailed statistical analysis related to **Figure 2**.

In the early groups, before the first enrichment period (pre-tests, future control and enriched groups pooled), mice did not discriminate Decanal and 2-Dodecanone, as shown by a similar investigation time between OHab4 and OTest (pre-tests, Fig. 2Bi and Table 2 for detailed statistics). After enrichment, the early-enriched group discriminated Decanal from 2-Dodecanone at 12, 15 and 18 months as evidenced by the increase of the investigation time between OHab4 and OTest, while control animals displayed no discrimination whatever their age (Fig. 2Bi and Table 2, with the exception of the early-control group at 12 months). Discrimination indices that allow evaluating the strength of the discrimination confirmed the better discrimination abilities of early-enriched mice compared to controls at 12, 15 and 18 months (Fig. 2Bii, Table 2). The late-enriched group also discriminated Decanal from 2-Dodecanone at 15 and 18 months while non-enriched controls did not (Fig. 2Ci, Table 2). Discrimination indices confirmed the higher discrimination abilities of late-enriched group compared to controls (Fig. 2Cii). The improvement of olfactory discrimination at 18 months in both early- and late-enriched aged mice was confirmed using another pair of perceptually similar odorants, (+) Limonene and (-) Limonene (Fig. 2Di-Eii).

All together, these results indicate that lifelong olfactory enrichment improves fine olfactory discrimination (*i.e.* discrimination of perceptually similar odorants) up to late ages. Importantly, this was true for both early and late-started enrichment.

### Long-run olfactory enrichment improves late age cognitive flexibility in a reversal task

In order to further investigate the cognitive benefits of long-run olfactory enrichment, mice were submitted to an olfactory associative learning paradigm followed by a rule reversal to assess cognitive flexibility (Fig. 3A).

In the early groups, before enrichment, young adult mice (2 months pre-tests, future enriched and controls pooled) displayed an increasing success rate with days of training (D1-D5), indicative of successful learning of the association of the odorant with the reward (Fig. 3Bi). At 12, 15 and 18 months, both early control and enriched animals similarly succeeded in this task (Fig. 3Bii-Biv, see Table 3 for detailed statistical results). Then, the learning rule was reversed (R1-R5). On R1, rule reversal induced a drop of performances reflecting the confidence of the animals in the previously learned rule. The drop in success rate was of the same magnitude for all ages of the early control and enriched groups (Fig. 3Bi-Biv, see Table 3 for detailed statistical results). From R1 to R5, early controls and enriched mice of 2, 12 and 15 months similarly learned the new rule indicating comparable behavioral flexibility (Fig. 3Bi-iii). A notable exception was the 18-month-old mice (Fig. 3Biv). Control animals learned the first rule (significant day effect and D5 > 50%), but upon reversal, the day effect did not reach significance and their performances were not different from chance level at R5, indicating unsuccessful reversal learning (Fig. 3Biv). In enriched animals, the day effect did not reach significance for the first rule (p=0.08), but performances at D5 were significantly different from chance level, suggesting that learning has occurred. For the reversal learning, enriched mice showed increase in performance with days of training (day effect p=0.012) and their performances exceeded chance level at R4 and R5, indicating successful reversal learning (Fig. 3Biv). This shed light on a benefit of an early starting long-run olfactory enrichment on cognitive flexibility at late age.

**Table 3:**
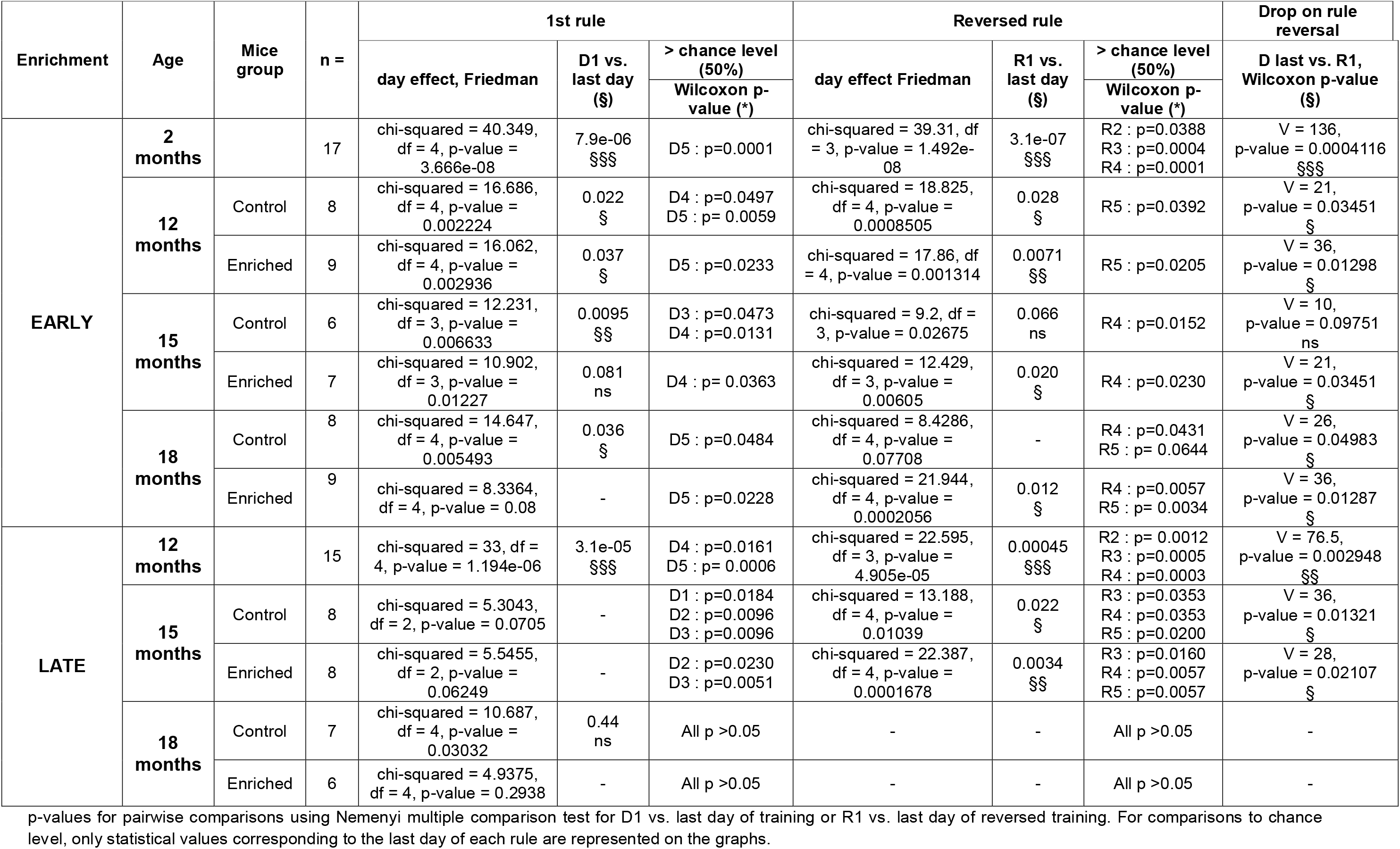
Detailed statistical analysis related to Figure 3.

In the late groups, before enrichment, 12-month-old animals (controls and enriched pooled) learned the associative task and the reversal task (Fig. 3Ci). At 15 months, controls and enriched animals similarly learned the associative task and the rule reversal (Fig. 3Cii). At 18 months, none of the groups (late-control or late-enriched) were able to learn the task as their success rate remained at chance level (50%) for 5 successive training days (Fig. 3Ciii), similar to what is usually observed for pseudo-conditioned groups (Supp. Fig. S2).

To sum up, long-run olfactory enrichment did not improve performances in a first associative learning but early-starting enrichment improved the ability to learn a second new associative rule (reversed rule), i.e. cognitive flexibility in late age.

### Long-run olfactory enrichment improves spatial memory at middle and late ages

In order to investigate whether long-run olfactory enrichment could benefit to a non-olfactory task, mice were submitted to the object location task to assess spatial memory (Fig. 4A).

In the early group, before any enrichment (pre-tests), young adult mice (2-month-old, future control and enriched pooled) detected the displaced object at all tested delays (5, 15 and 60 min; Fig. 4Bi, see Table 4 for detailed statistical results), as shown by a preference index that was significantly higher than the chance level (50%). At middle age (12 months), early-control mice showed a retention deficit at 60 min, while, on the contrary, 12-month-old early-enriched mice were able to remember the object location after 60 min (Fig. 4Bii). This result indicated that long-run olfactory enrichment allowed maintaining spatial memory performances. At 15 months, early-control mice were able to remember the object location for the 5- and 15-min delays but not for the 60-min delay (Fig. 4Biii), a pattern similar to that of early-control mice aged 12 months. This result indicated a stability of the spatial memory performances between 12 and 15 months. No benefits of early enrichment were visible at 15 months (Fig. 4Biii). At 18 months, early-control mice were unable to remember the object location whatever the delay (Fig. 4Biv), indicating a further reduction of the retention time of spatial information, compared to 12 or 15-month-old animals. In contrast, early-enriched 18-month-old mice succeeded in detecting the displaced object for all tested delays (Fig. 4Biv), revealing a better ability of 18-month-old mice to retain spatial information when submitted to long-run olfactory enrichment In late groups, mice pre-tested at 12 months (future control and enriched pooled) showed retention of the object location at 5 and 15 min but not at 60 min (Fig 4Ci), replicating the data presented on Fig. 4Bii with another group of animals. At 15 months, late-enriched mice but not late-control mice displayed a preference for the displaced object at the 15-min delay (Fig. 4Cii). This result demonstrated that the late-started olfactory enrichment efficiently lengthened the retention of spatial information up to 15 min. At 18 months (Fig. 4Ciii), late-controls showed retention of the object location at 15 min and 60 min but not 5 min. Early- and late-controls thus showed some discrepancies at the 15- and 60-min delays, suggesting some instability of performances from one group of animals to another in the late ages. However, it is worth noting that the late-enriched mice retained spatial information up to 15 min, similarly to early-enriched mice. These results suggested that the late-started enrichment still had benefits on the retention of spatial memory.

**Table 4:** Detailed statistical analysis related to Figure 4.

| Enrichment | Age | Mice group | 5 min |  | 15 min |  | 60 min |  |
| --- | --- | --- | --- | --- | --- | --- | --- | --- |
|  |  |  | n = | Wilcoxon, p-value | n = | Wilcoxon, p-value | n = | Wilcoxon, p-value |
| EARLY | 2 months |  | 29 | V = 322, p-value = 0.01145<br>* | 30 | V = 334, p-value = 0.006066<br>** | 28 | V = 405, p-value = 2.231e-06<br>*** |
|  | 12 months | Control | 15 | V = 120, p-value = 3.052e-05<br>*** | 15 | V = 111, p-value = 0.001007<br>** | 15 | V = 87, p-value = 0.06769<br>n.s. |
|  |  | Enriched | 15 | V = 119, p-value = 6.104e-05<br>*** | 17 | V = 149, p-value = 5.341e-05<br>*** | 17 | V = 129, p-value = 0.005493<br>** |
|  | 15 months | Control | 12 | V = 68, p-value = 0.0105<br>* | 14 | V = 89, p-value = 0.01013<br>* | 14 | V = 67, p-value = 0.1897<br>n.s. |
|  |  | Enriched | 14 | V = 104, p-value = 0.0006815<br>*** | 15 | V = 95, p-value = 0.02396<br>* | 16 | V = 76, p-value = 0.3529<br>n.s. |
|  | 18 months | Control | 12 | V = 52, p-value = 0.3424<br>n.s. | 13 | V = 68, p-value = 0.0636<br>n.s. | 12 | V = 54, p-value = 0.1273<br>n.s. |
|  |  | Enriched | 12 | V = 78, p-value = 0.0002441<br>*** | 14 | V = 105, p-value = 6.104e-05<br>*** | 15 | V = 107, p-value = 0.004107<br>** |
| LATE | 12 months |  | 32 | V = 521, p-value = 4.424e-09<br>*** | 30 | V = 441, p-value = 9.415e-06<br>*** | 32 | V = 260, p-value = 0.5335<br>n.s. |
|  | 15 months | Control | 15 | V = 109, p-value = 0.001678<br>** | 11 | V = 41, p-value = 0.2598<br>n.s. | 10 | V = 25, p-value = 0.4055<br>n.s. |
|  |  | Enriched | 13 | V = 86, p-value = 0.001221<br>** | 14 | V = 96, p-value = 0.002014<br>** | 13 | V = 32, p-value = 0.5531<br>n.s. |
|  | 18 months | Control | 12 | V = 42, p-value = 0.425<br>n.s. | 13 | V = 82, p-value = 0.004028<br>** | 12 | V = 61, p-value = 0.04614<br>* |
|  |  | Enriched | 14 | V = 104, p-value = 0.0001221<br>*** | 12 | V = 74, p-value = 0.001709<br>** | 10 | V = 36, p-value = 0.2158<br>n.s. |

Univariate Wilcoxon tests were used to compare the preference index to 50%.

Thus, spatial memory is degraded at middle age (12-15 months) with decreased retention time that globally aggravated with further aging. Long-run olfactory enrichment is beneficial, whether started in young adulthood or at middle age, with a somehow more pronounced effect in the early-enriched group.

To conclude on the behavioral evaluations, long-run olfactory enrichment proved efficient, both when started in young adulthood or at middle-age, in increasing olfactory discrimination and higher order cognitive processes including flexibility evaluated in an olfactory associative task, and spatial memory. The benefits of long-run olfactory enrichment thus extend outside of the olfactory sphere. This finding is further supported by a significant positive correlation between spatial memory (5-min delay) and olfactory discrimination performances (ρ(45)=0.5519849, p=5.132e-05) at 18 months (Fig. 4D), suggesting an overall positive effect of long-run olfactory enrichment on cognition.

### Long-run olfactory enrichment induces remodeling of NA innervation

In search for mechanisms that could support the cognitive benefits of long-run olfactory enrichment at the cellular level, we focused on the noradrenergic system. We hypothesized that long-run olfactory enrichment would activate the noradrenergic system and thereby would preserve this neuromodulatory system from aging, as could be predicted based on the noradrenergic theory of the cognitive reserve (Robertson, 2013; Clewett et al., 2016; Mather and Harley, 2016). We thus investigated the integrity of the *Locus Coeruleus* and the density of NA innervation in the 18-month-old mice after early or late start long-run olfactory enrichment.

Using immunohistochemistry of the transcription factor Phox2a specifically expressed in noradrenergic neurons (Morin et al., 1997; Yang et al., 1998; Goridis and Brunet, 1999) (Fig. 5A), we measured the surface of the *Locus Coeruleus* (data not shown) and the density of noradrenergic neurons (Fig. 5Bi). We did not observe any effect of the onset of enrichment and no interaction with enrichment (2-way ANOVA, see Table 5 for detailed statistical results) and we pooled the early- and late-enriched groups, as well as their respective control groups. Control and enriched mice displayed similar densities of Phox2a-positive neurons (Fig. 5Bi, p>0.05). Similar densities of Tyrosine Hydroxylase positive cells (TH, enzyme required for NA synthesis and marker of NA neurons) were also found in the *Locus Coeruleus* of enriched and control mice (Fig. 5A and Bii, see Table 5 for detailed statistical results). Moreover, “resting-noradrenergic” neurons exist in the rostral *Locus Coeruleus* under basal conditions. These neurons express the Phox2a transcription factor but not TH and can be reactivated following treatment with RU 24722 (a catecholaminergic activator) (Bezin et al., 2000). The percentage of Phox2a-positive cells co-expressing TH was about 95% in both groups, indicating that there was no reactivation of TH expression in Phox2a-positive cells upon long-run olfactory enrichment (Fig. 5Biii, see Table 5 for detailed statistical results). Finally, as TH expression can be graded, we assessed the level of TH expression in noradrenergic neurons that could be modified by long-run olfactory enrichment. We measured the optical density of TH staining in the *Locus Coeruleus* and found no difference in enriched animals compared to controls (Fig. 5C, see Table 5 for detailed statistical results). Altogether, these results indicated that long-run olfactory enrichment did not change the density of noradrenergic neurons in the *Locus Coeruleus* of aged mice nor NA synthesis machinery.

**Table 5:** Detailed statistical analysis related to Figure 5.

|  | n= -mice per group | 2-way ANOVA |
| --- | --- | --- |
| Phox2A+ cell density | n=6 for early control, n=6 for early enriched, n=4 for late control, n=4 for late enriched | enrichment effect $F_{(1,16)}=0.6438$, $p=0.4341$<br>onset of the enrichment $F_{(1,16)}=0.0295$, $p=0.8658$<br>interaction enrichment x onset $F_{(1,16)}=0.0779$, $p=0.7837$ |
| TH+ cell density | n=6 for early control, n=6 for early enriched, n=4 for late control, n=4 for late enriched | enrichment effect $F_{(1,16)}=1.0850$, $p=0.3131$<br>onset of the enrichment $F_{(1,16)}=0.0005$, $p=0.9818$<br>interaction enrichment x onset $F_{(1,16)}=0.1809$, $p=0.6763$ |
| Phox2A+/TH+ cell percentage | n=6 for early control, n=6 for early enriched, n=4 for late control, n=4 for late enriched | enrichment effect $F_{(1,16)}=1.4386$, $p=0.2478$<br>onset of the enrichment $F_{(1,16)}=1.6375$, $p=0.2189$<br>interaction enrichment x onset $F_{(1,16)}=0.3852$, $p=0.5436$ |
| TH intensity | n=6 for early control, n=6 for early enriched, n=4 for late control, n=5 for late enriched | enrichment effect $F_{(1,16)}=1.2485$, $p=0.27938$<br>onset of the enrichment $F_{(1,16)}=3.7019$, $p=0.07126$<br>interaction enrichment x onset $F_{(1,16)}=0.0421$, $p=0.83977$ |

After pooling early- and late mice
|  | n= mice per group | Unilateral unpaired Student t-test or unilateral unpaired Wilcoxon test |
| --- | --- | --- |
| Phox2A+ cell density | n= 10 for control group, n= 10 for enriched group | t = -0.85407, df = 18, p-value = 0.2021 |
| TH+ cell density | n= 10 for control group, n= 10 for enriched group | W = 37, p-value = 0.1763 |
| Phox2A+/TH+ cell percentage | n= 10 for control group, n= 10 for enriched group | t = -1.041, df = 18, p-value = 0.1558 |
| TH intensity | n= 10 for control group, n= 11 for enriched group | t = -1.1394, df = 19, p-value = 0.1343 |

**Figure 5:**
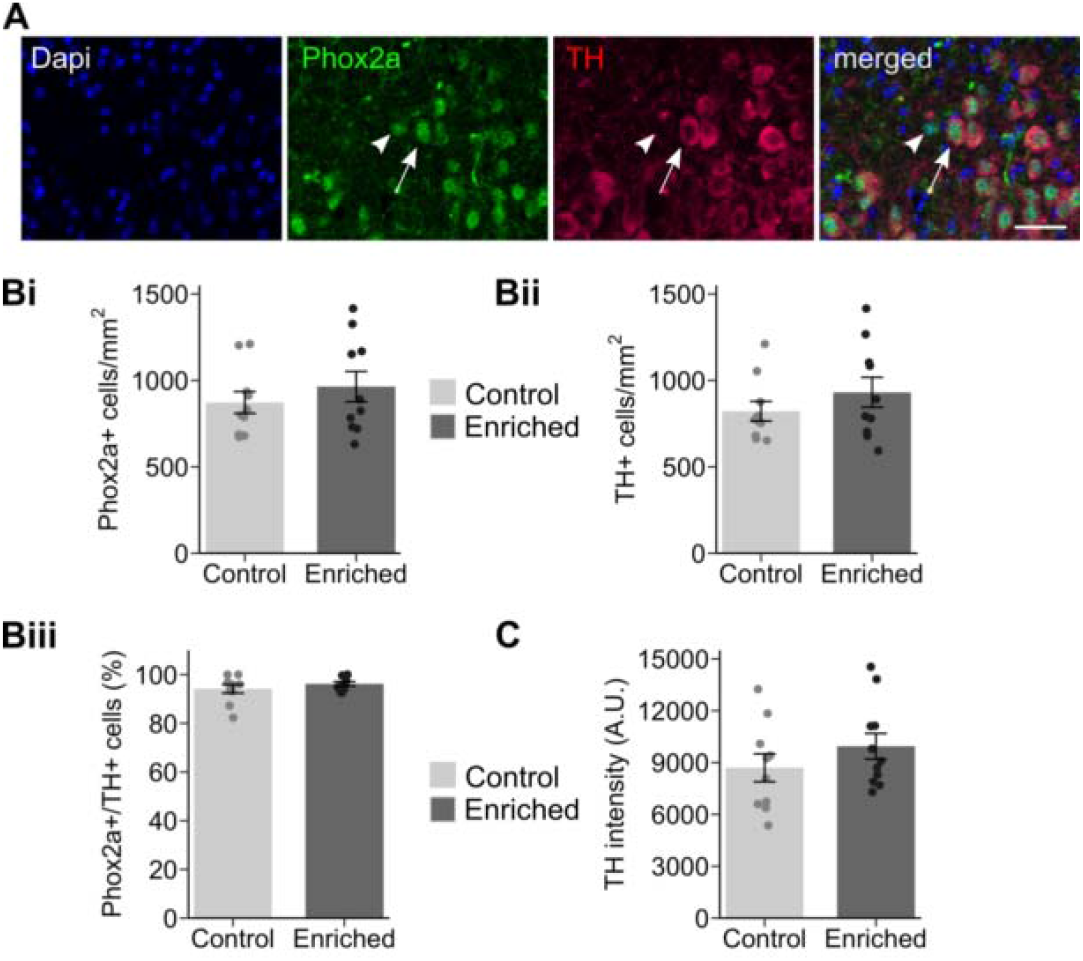
No impact of long-run olfactory enrichment on the integrity of the *Locus Coeruleus*. Phox2a and TH double-immunolabeling allowed to analyze the integrity of the *Locus Coeruleus*. **(A)** Example of a Phox2a+/TH-neuron (arrowhead) and a Phox2a+/TH+ neuron (arrow). **(B-C)** Quantification of Phox2a+ cell density **(Bi)**, TH+ cell density **(Bii)**, Phox2a+/TH+ double-labeled cells **(Biii)** and TH intensity **(C)** in the *Locus Coeruleus* revealed no difference between enriched and control mice s. Data represent the mean ± SEM. Dots represent data from individual mice. n=10-11 mice per group. Scale bar = 40µm.

In order to quantify noradrenergic innervation, we then evaluated the density of NET-positive fibers (NorEpinephrine Transporter) in brain regions related to the tested behavioral performances: the olfactory bulb for olfactory discrimination (*e.g.* Moreno et al., 2009, 2014; Rey et al., 2012; Huang et al., 2016) (Fig. 6Ai), the dorsal hippocampus for spatial memory (*e.g.* Moser et al., 1995; Assini et al., 2009; Fanselow and Dong, 2010; Eichenbaum, 2017) (Fig. 6Aii) and the lateral orbitofrontal cortex (LO) for cognitive flexibility (*e.g.* Schoenbaum et al., 2002; Ragozzino, 2007; Izquierdo et al., 2017) (Fig. 6Aiii). More precisely, the density of NET-positive fibers was evaluated in the granule cell layer of the olfactory bulb (where it is the highest (Shipley et al., 1985; McLean et al., 1989)), the CA1 (stratum oriens, pyramidal cell layer and stratum radiatum), DG (granule-cell and molecular layers) and CA3 regions of the dorsal hippocampus, and the lateral orbitofrontal cortex in enriched and control mice. As we did not observe any difference between the early and late groups (p>0.05 for all comparisons, see Table 6 for detailed statistics), we pooled the early- and late-enriched groups, as well as their respective control groups. In the olfactory bulb, the density of NA fibers was significantly increased (+ 23%) in enriched mice compared to controls (Fig. 6Bi, unilateral Wilcoxon test, W=29, p=0.021). In the CA1 of the dorsal hippocampus, the density of NA fibers was also significantly higher (+45%) in enriched mice compared to controls (Fig. 6Bii, W=34, p=0.047). This increase was not observed in the DG (Fig. 6Bii, unilateral t.test, t=-0.39997, p=0.3467), CA3 (Fig. 6Bii, t = 1.319, df = 8, p-value = 0.2237) and in the lateral orbital cortex (Fig. 6Biii, W = 34, p-value = 0.1527). Finally, the NA fiber density in CA1 was positively correlated with spatial memory performances at the 5-min delay (Fig. 6Ci, ρ (20)=0.3894737, p=0.04529). In the same way, in the olfactory bulb, the NA innervation was positively associated with olfactory discrimination performances (Fig. 6Cii, ρ(21)=0.3657032, marginal significance p=0.05152).

**Table 6:**
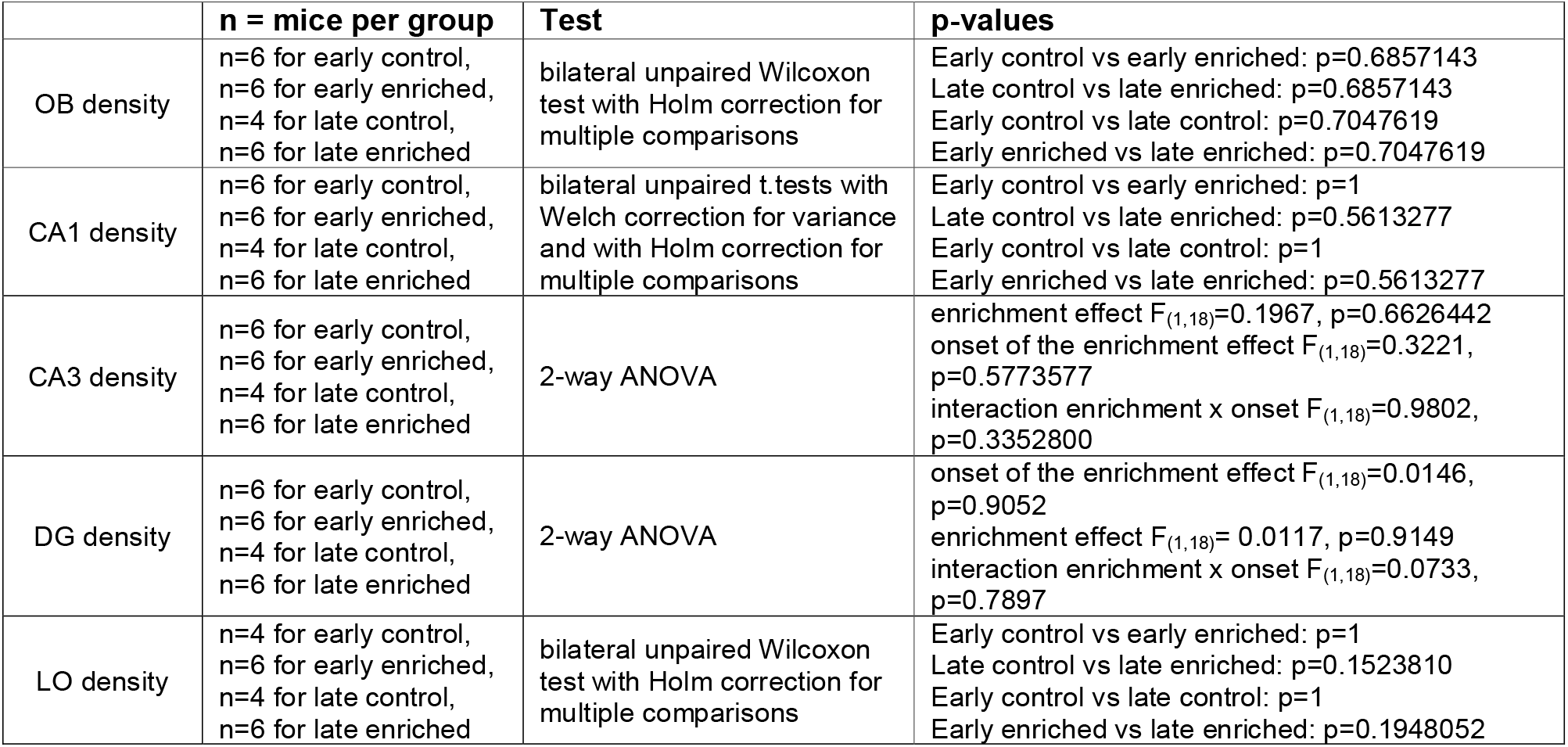
Detailed statistical analysis related to **Figure 6**.

|  | n = mice per group | Test | p-values |
| --- | --- | --- | --- |
| OB density | n=6 for early control,<br>n=6 for early enriched,<br>n=4 for late control,<br>n=6 for late enriched | bilateral unpaired Wilcoxon test with Holm correction for multiple comparisons | Early control vs early enriched: p=0.6857143<br>Late control vs late enriched: p=0.6857143<br>Early control vs late control: p=0.7047619<br>Early enriched vs late enriched: p=0.7047619 |
| CA1 density | n=6 for early control,<br>n=6 for early enriched,<br>n=4 for late control,<br>n=6 for late enriched | bilateral unpaired t.tests with Welch correction for variance and with Holm correction for multiple comparisons | Early control vs early enriched: p=1<br>Late control vs late enriched: p=0.5613277<br>Early control vs late control: p=1<br>Early enriched vs late enriched: p=0.5613277 |
| CA3 density | n=6 for early control,<br>n=6 for early enriched,<br>n=4 for late control,<br>n=6 for late enriched | 2-way ANOVA | enrichment effect $F_{(1,18)}=0.1967$, p=0.6626442<br>onset of the enrichment effect $F_{(1,18)}=0.3221$, p=0.5773577<br>interaction enrichment x onset $F_{(1,18)}=0.9802$, p=0.3352800 |
| DG density | n=6 for early control,<br>n=6 for early enriched,<br>n=4 for late control,<br>n=6 for late enriched | 2-way ANOVA | onset of the enrichment effect $F_{(1,18)}=0.0146$, p=0.9052<br>enrichment effect $F_{(1,18)}=0.0117$, p=0.9149<br>interaction enrichment x onset $F_{(1,18)}=0.0733$, p=0.7897 |
| LO density | n=4 for early control,<br>n=6 for early enriched,<br>n=4 for late control,<br>n=6 for late enriched | bilateral unpaired Wilcoxon test with Holm correction for multiple comparisons | Early control vs early enriched: p=1<br>Late control vs late enriched: p=0.1523810<br>Early control vs late control: p=1<br>Early enriched vs late enriched: p=0.1948052 |

**Figure 6:**
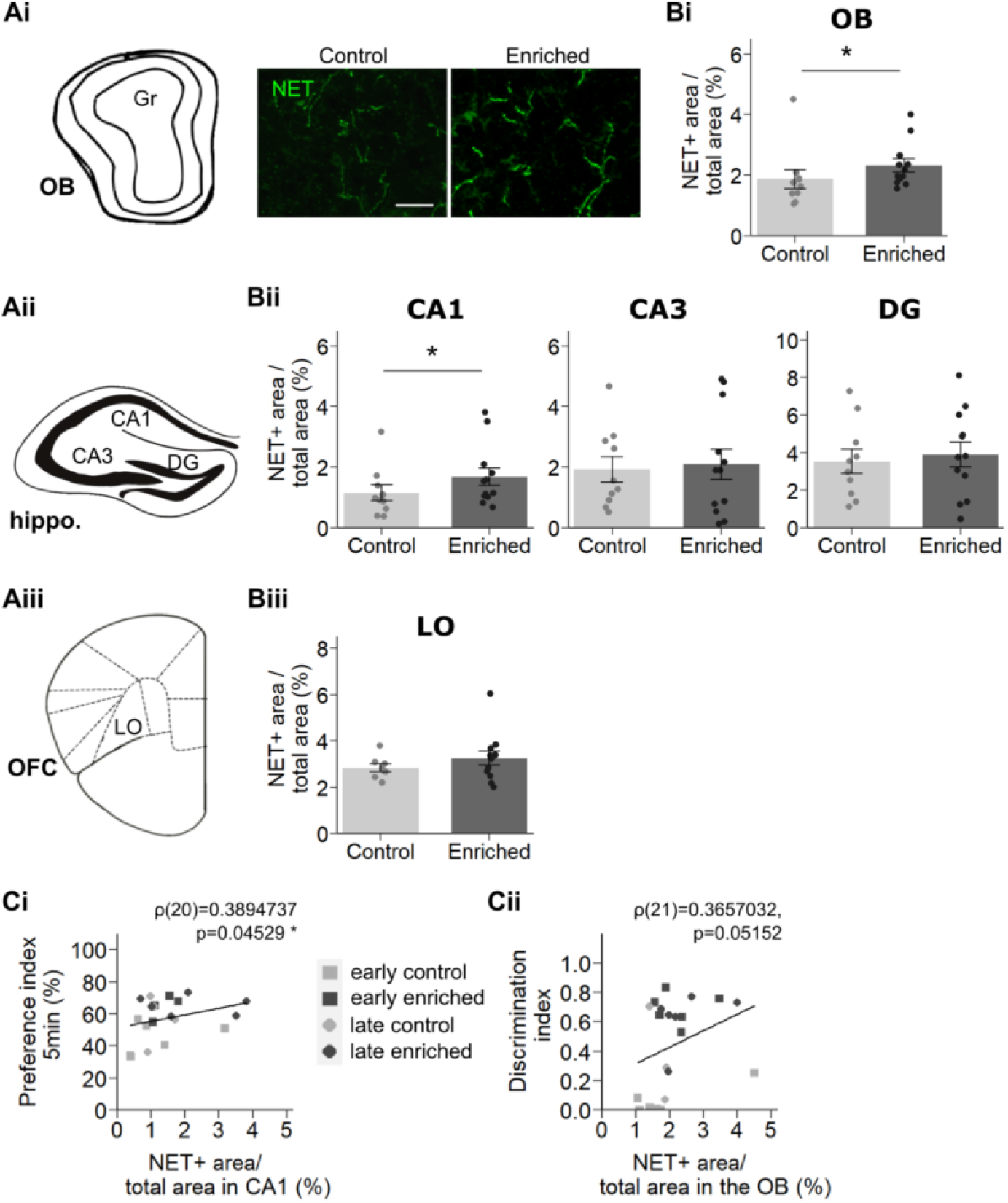
Local increases of noradrenergic innervation in the aged brain following long-run olfactory enrichment positively correlate with behavioral performances. **(A)** Norepinephrine transporter (NET) immunolabeling allowed to visualize NA fibers within the granule cell layer of the olfactory bulb **(Ai)**, the hippocampus **(Aii)** and the OFC **(Aiii)**. **(B)** Quantification of the area occupied by NET-positive fibers in the olfactory bulb **(Bi)**, CA1, dentate gyrus and CA3 of the hippocampus **(Bii)** and LO **(Biii)** of enriched and control mice. Data represent the mean ± SEM. Dots represent data from individual mice. n=8-12 mice per group. **(C)** Significant positive linear correlations were found between the NA innervation in the dorsal hippocampal CA1 and the preference index for the 5-min-delay **(Ci)** and between the NA innervation in the olfactory bulb and the discrimination index for the Decanal / Dodecanone pair **(Cii)**. n= 20-21 mice. *p<0.05. DG: Dentate Gyrus; Gr: granule cell layer of the olfactory bulb; LO: Lateral Orbital cortex, OB: Olfactory Bulb, OFC: OrbitoFrontal Cortex. Scale bar = 20µm

In summary, our results revealed an increase of noradrenergic innervation in the olfactory bulb and CA1 following long-run olfactory enrichment in 18-month-old enriched mice that is linked with the behavioral improvements.

### Long-run olfactory enrichment induced task-specific changes in brain functional connectivity

As preserved cognitive functions are often associated with remodeling of brain activation patterns in aged humans (Grady, 2012; Cabeza et al., 2018), we hypothesized that long-run olfactory enrichment drives changes in brain activation levels or/and functional connectivity that could contribute to improvement in behavioral performances. Brain activation was investigated in controls and enriched mice by *post-mortem* mapping of the expression of the immediate early gene c-Fos (Midroit et al., 2018), following the olfactory discrimination task or the object location task. C-Fos-positive cell mapping allows quantification and statistical analysis of task-evoked neuronal activation and connectivity in multiple brain regions. This analysis was performed on pooled early and late groups for the olfactory discrimination task (n=7 control: 5 early + 2 late, and n=7 enriched mice: 5 early + 2 late). and on early groups only for the object location task with 60-min delay (n=5 control and n=5 enriched mice).

The density of activated neurons was measured in 31 brain regions (Supp. Fig. S3). To contrast differences between enriched and control animals, we expressed c-Fos density as a percentage of control mice. Results showed that enriched animals had an activation level similar to that of controls for both tasks (Fig.7 and Table 7), except for a lower activation of the frontal region (Fr) in enriched animals for the olfactory discrimination task (Fig.7A, bilateral one sample t.test corrected for multiple comparisons, t=-6.4513016, p=0.020367). This first analysis revealed that long-run olfactory enrichment did not induce major changes in terms of density of neurons activated by the tasks.

**Table 7:**
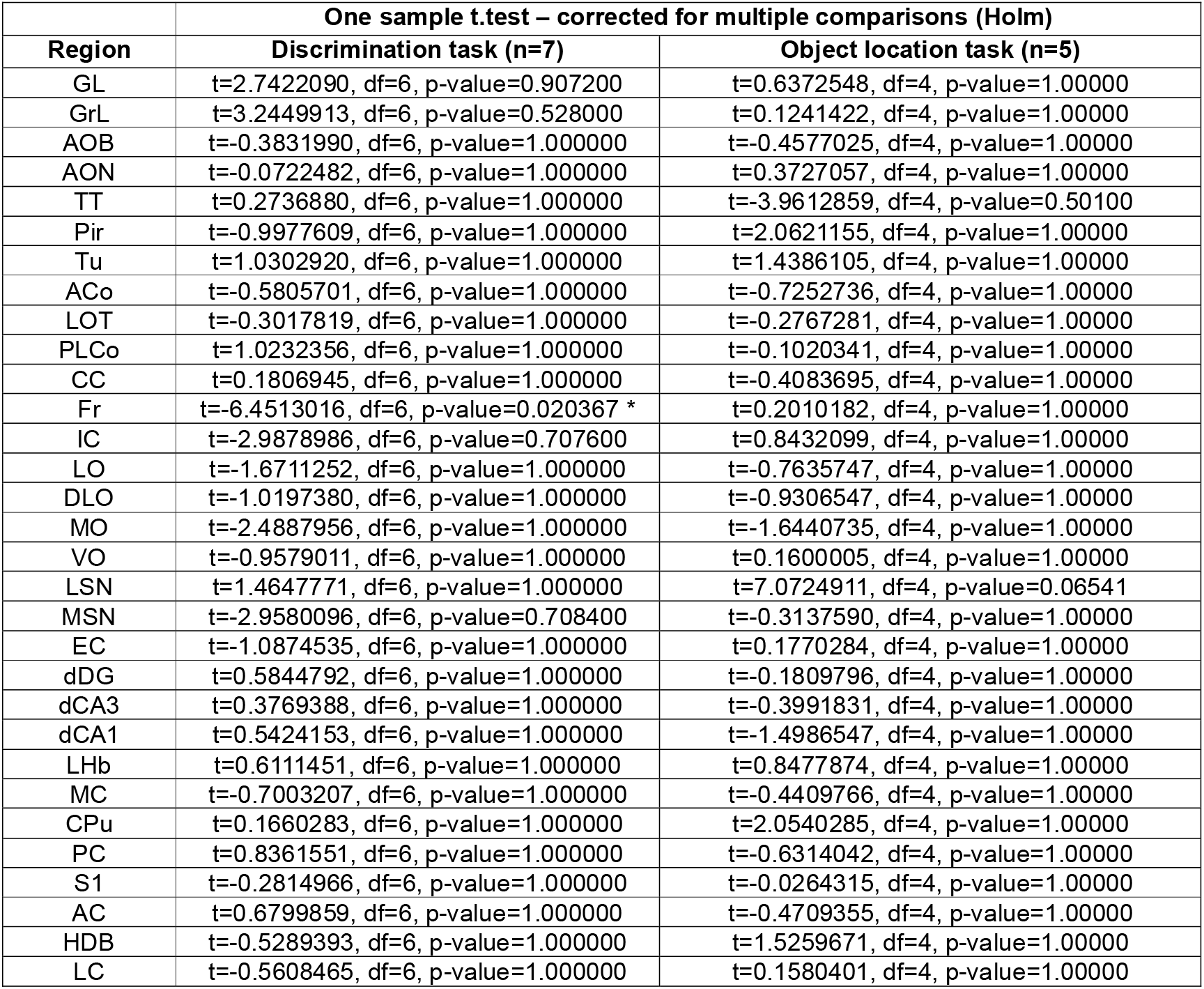
Detailed statistical analysis related to **Figure 7**.

**Figure 7:**
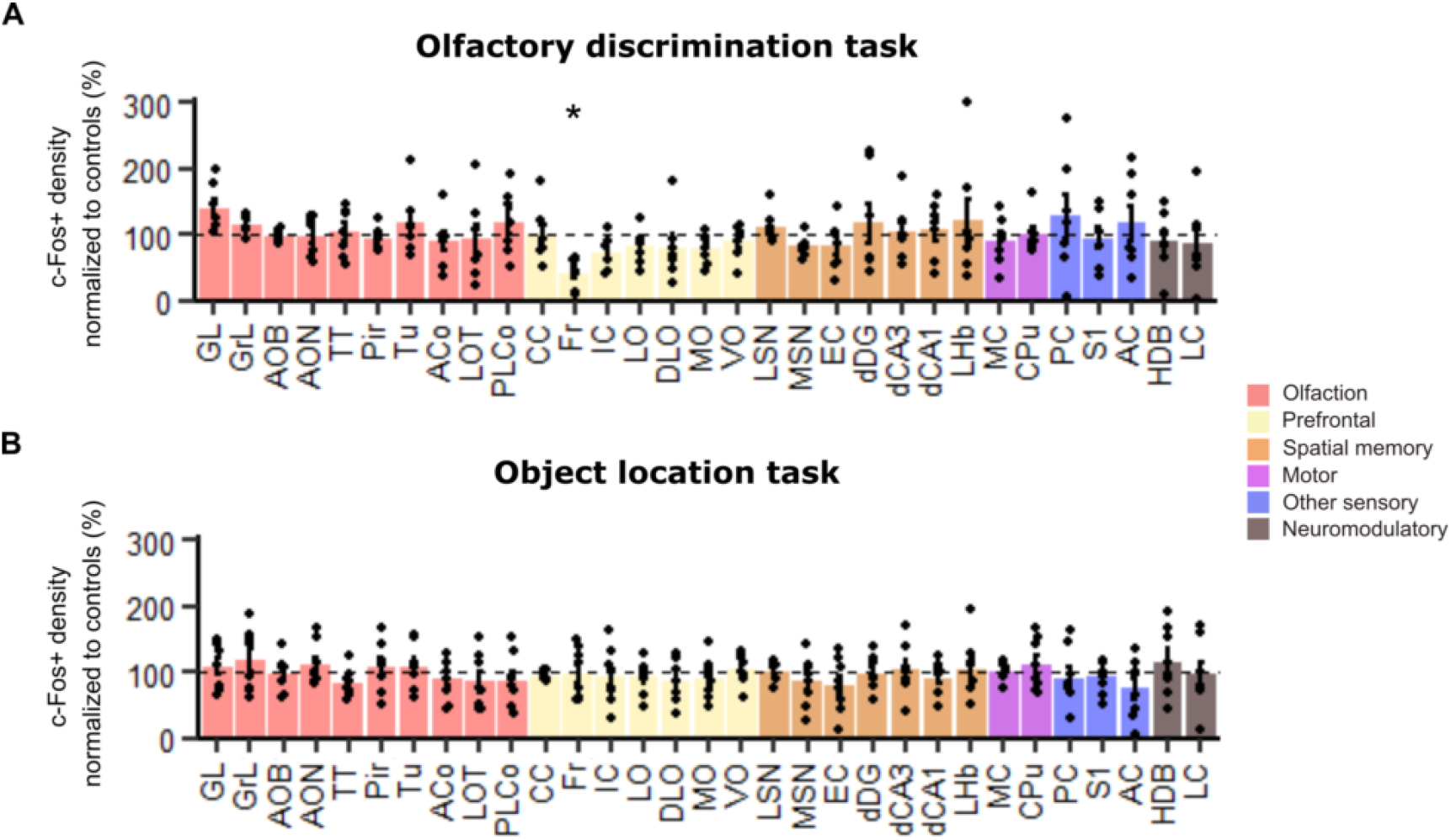
Brain regional activation induced by the olfactory discrimination task and the object location task. c-Fos+ cell densities of enriched mice for the 31 regions of interest in the olfactory discrimination task **(A)** and the object location task **(B)**. Results of enriched mice are expressed as percentages of controls. Data represent the mean ± SEM. Dots represent data from individual mice. n=5-7 mice per group. Statistical comparisons to 100% were assessed by bilateral one-sample t.test corrected for multiple comparisons. *p<0.05.

We then investigated functional connectivity between the 31 brain regions included in the analysis. For that purpose, we calculated correlations in c-Fos-positive cell density between all the pairs of brain regions within each group (olfactory discrimination task: controls and enriched, object location task: controls and enriched, Fig.8A and D) (Wheeler et al., 2013; Babayan et al., 2017). Correlations between structures, reflecting functional connections, were thresholded (Pearson’s r ≥ 0.82, corresponding to a bilateral significance level of p ≤ 0.025) (Fig.8B and E). For both tasks, different thresholds were tested and revealed the same overall trends as those described below (Supp. Fig. 4).

**Figure 8:**
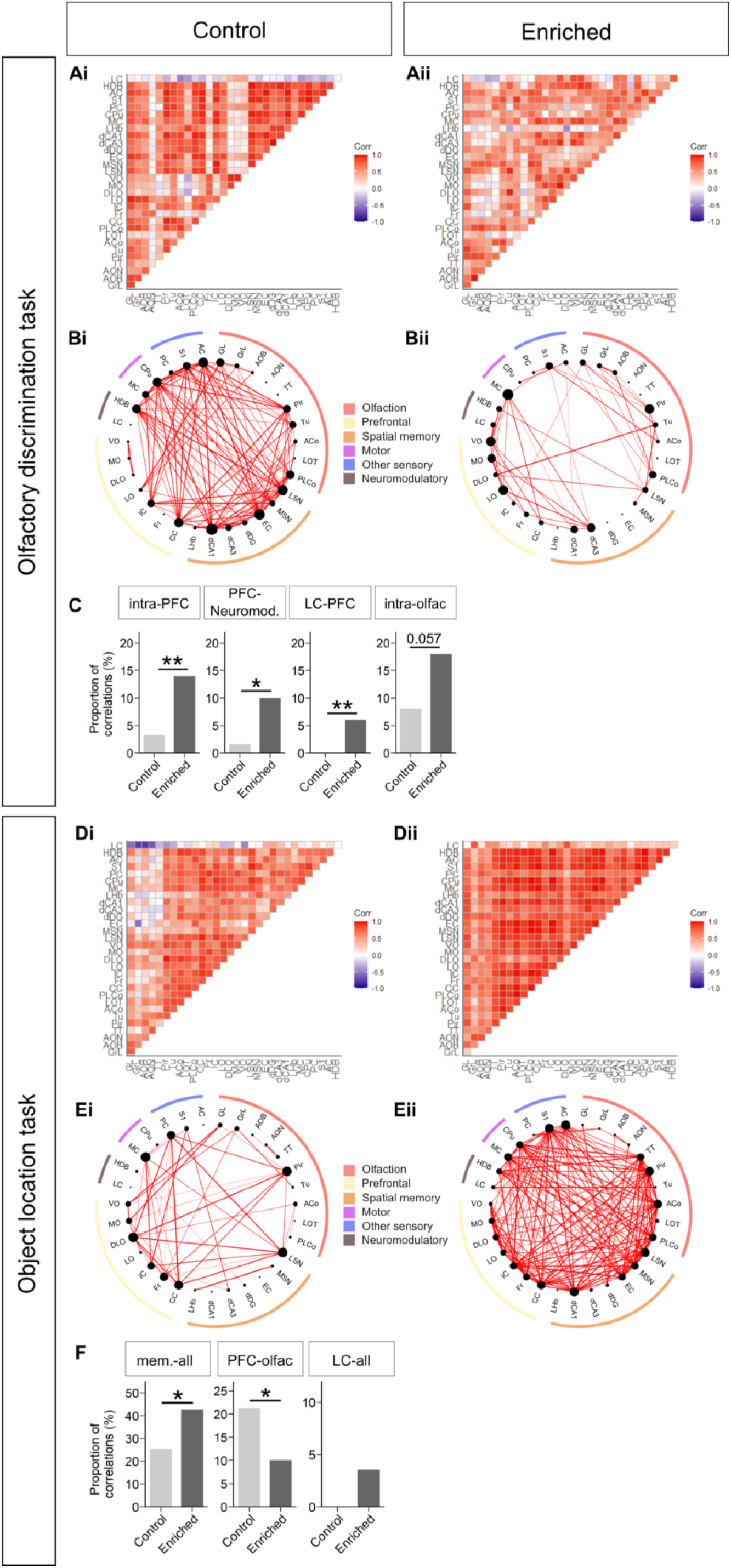
Task-dependent reorganization of functional networks following long-run olfactory enrichment. Functional networks were analyzed in controls (left column) and enriched animals (right column) in the olfactory discrimination task **(A-C)** and the object location task **(D-F)**. **(Ai-Aii, Di-Dii)** Matrices representing all inter-regional correlations (without any thresholding). Axes correspond to brain structures and colors reflect correlation strength (with positive correlations in red and negative correlations in blue). **(Bi-Bii, Ei-Eii)** Networks representing correlations between regions. Graphs were generated by considering only the strongest correlations (Pearson’s r ≥ 0.82, p ≤ 0.025). The thickness of the connection line is proportional to the correlation strength and the node size is proportional to the node’s degree (number of connections). **(C and F)** Graphics representing the percentage of connections within or between systems among the effective (significant) correlations in each group. Statistical analyses were assessed by bilateral proportion tests. *p<0.05, **p<0.01.

Regarding the olfactory discrimination task, network analysis revealed that enriched mice displayed less connections than controls (Fig.8Bi-Bii, see Table 7 for detailed statistics). Indeed, the density of connections in enriched mice decreased by 60% when compared to control mice (controls: 124 connections, enriched: 50 connections, out of 465 possible, proportion test X-squared = 38.715, p=4.905e-10). The reduction in the density of connections affected all functional systems, except the prefrontal and neuromodulatory systems in which it remained unchanged (see Table 7 for detailed connections counts and statistics). Beside the overall change in the connection density, we analyzed the relative load of each brain functional system among the effective (significant) connections. This analysis indicated an increase of the load of the PFC connections within the network of enriched mice. Indeed, the proportion of connections between regions belonging to the PFC system (intra-PFC) and the connections between the PFC system and the neuromodulatory system increased compared to controls (Fig.8C, controls: 4 connections out of 124 effective, enriched: 7 out of 50, X-squared = 6.9844, p=0.008222 for intra-PFC, controls: 2 connections out of 124 effective, enriched: 5 out of 50, X-squared = 6.4916, df = 1, p-value = 0.01084 for PFC-Neuromodulatory). In addition, the *Locus Coeruleus* was connected with prefrontal structures (LC-PFC) in enriched mice, while it did not correlate with any structures in control mice (Fig.8C, controls: 0 connection out of 124 effective, enriched: 3 out of 50, X-squared = 7.5705, p=0.005933). The connections between olfactory regions (intra-olfac) also increased in proportion with enrichment, reaching a marginal statistical significance (Fig. 8BC, controls: 10 connections out of 124 effective, enriched: 9 out of 50, X-squared = 3.616, p= 0.05722). This network analysis suggested that lifelong olfactory enrichment induced a reorganization of the brain network associated with the olfactory discrimination task with a global reduction of connections but an increase in the relative contribution of the PFC and the olfactory system to the functional network.

To go further into the analysis of these networks’ reorganization and describe the networks’ topology, we performed a graph theory analysis (Rubinov and Sporns, 2010; Wheeler et al., 2013; Babayan et al., 2017). We measured efficiency and clustering coefficients as indexes of functional integration and segregation respectively. The networks activated by the olfactory discrimination task were segregated as they displayed a higher clustering coefficient compared to random networks for both control and enriched groups (Supp. Fig. S5Ai-Aii). The level of both control and enriched networks’ integration were equivalent to random networks (Supp. Fig. S5Bi-Bii). Thus, the functional networks engaged by the olfactory discrimination task displayed properties that were consistent with a small-world topology. We pursued the analyses by applying a Markov clustering algorithm, which organizes structures into clusters based on their common inter-connections. Two clusters were identified in the control group performing the olfactory discrimination task versus 4 in the enriched group (Supp. Fig. S5Ci-Cii). This difference in clustering between control and enriched mice added support to the notion that the network underlying olfactory discrimination became more segregated after long-run olfactory enrichment. Combined with the non-homogeneous decrease in functional connections described above, these findings suggested that enriched animals displayed a more specialized network for the olfactory discrimination task than controls.

In contrast to the olfactory discrimination task, the density of connections in the object location task, was significantly higher in enriched mice than in control mice (Fig.8Ei-Eii, controls: 47 connections, enriched: 169 connections out of 465 possible, X-squared = 89.753, p<2.2e-16) reaching a 260% increase. The increase in the density of connections affected all functional systems (see Table 8 for detailed connections counts and statistics). However, looking at the relative contribution of each system with regard to the effective connections, spatial memory regions turned out to be more connected with the rest of the brain (mem.-all) in enriched mice compared to controls (Fig.8F, controls: 12 connections out of 47 effective, enriched: 72 out of 169, X-squared = 4.5096, p=0.03371) with an increase of 67%. Notably, entorhinal cortex (EC) and dCA1 were not connected in controls but displayed connections in enriched animals (Fig8Ei-Eii), leading to a statistically significant increase of their contribution to the network (controls: 0 connection out of 47 effective, enriched: 15 out of 169, X-squared = 4.4829, df = 1, p-value = 0.03424 for EC; controls: 0 connection out of 47 effective, enriched: 19 out of 169 X-squared = 5.7937, df = 1, p-value = 0.01608 for dCA1). In addition, although the PFC system was globally more connected in enriched animals compared to controls, its connectivity increased less than that of other brain systems and its relative load in the network decreased, with a lower connection density between the PFC and the olfactory system (controls: 10 connections out of 47 effective, enriched: 17 out of 169, X-squared = 4.2306, df = 1, p-value = 0.0397). Last, the *Locus Coeruleus* showed no connection with any structures (LC-all) in control mice whereas it was connected with 6 structures in enriched mice, although this change was not statistically significant (Fig.8F, X-squared = 1.7163, df = 1, p-value = 0.1902). Connectivity of the auditory cortex also increased (controls: 0 connection out of 47 effective, enriched: 20 out of 169, X-squared = 6.1297, df = 1, p-value = 0.01329). The networks activated by the object location task displayed a clustering coefficient similar to that of random networks for both control and enriched groups (Supp. Fig. S6). Thus, the organization of the functional networks engaged by the object location task were not consistent with a small-world topology and we did not perform the cluster analysis. To sum up, brain activation associated with the object location task was reorganized in enriched mice with an increased density of connections, including new connections involving entorhinal cortex, dorsal CA1 and the *Locus Coeruleus*.

**Table 8:** Detailed number of correlations and proportions tests for the olfactory discrimination task, related to Figure 8.

|  | OLFACTORY DISCRIMINATION TASK |  |  |  |  |
| --- | --- | --- | --- | --- | --- |
|  | Number of correlations |  | Proportion of correlations (ratio of theoretical possible correlations) |  |  |
|  | Control | Enriched | Control | Enriched | Comparison Control versus Enriched |
| Total_theoretical_correlations | 465 | 465 |  |  |  |
| Total_effective_correlations | 124 | 50 | 0.2666667 | 0.1075269 | **X-squared = 38.715, df = 1, p-value = 4.905e-10 \*\*\*** |
| Olfaction_all | 53 | 18 | 0.11397849 | 0.03870968 | **X-squared = 18.68, df = 1, p-value = 1.546e-05 \*\*\*** |
| PFC_all | 31 | 24 | 0.06666667 | 0.05161290 | X-squared = 0.94691, df = 1, p-value = 0.3305 |
| Memory_all | 65 | 16 | 0.1397849 | 0.0344086 | **X-squared = 32.47, df = 1, p-value = 1.21e-08 \*\*\*** |
| Motor_all | 23 | 9 | 0.04946237 | 0.01935484 | **X-squared = 6.3433, df = 1, p-value = 0.01178 \*** |
| Sensory_all | 32 | 8 | 0.0688172 | 0.0172043 | **X-squared = 15.047, df = 1, p-value = 0.0001049 \*\*\*** |
| Neuromodulators_all | 14 | 7 | 0.03010753 | 0.01505376 | X-squared = 2.3872, df = 1, p-value = 0.1223 |

|  |  | Number of correlations |  | Proportion of correlations (ratio of effective correlations) |  |  |
| --- | --- | --- | --- | --- | --- | --- |
|  |  | Control | Enriched | Control | Enriched | Comparison Control versus Enriched |
|  | Total_effective_correlations | 124 | 50 |  |  |  |
| OLFACTION | Olfaction_all | 53 | 18 | 0.4274194 | 0.3600000 | X-squared = 0.67052, df = 1, p-value = 0.4129 |
|  | Intra_Olfaction | 10 | 9 | 0.08064516 | 0.18000000 | **X-squared = 3.616, df = 1, p-value = 0.05722 (#)** |
|  | Olfaction_PFC | 8 | 2 | 0.06451613 | 0.04000000 | X-squared = 0.39537, df = 1, p-value = 0.5295 |
|  | Olfaction_Memory | 19 | 4 | 0.1532258 | 0.0800000 | X-squared = 1.6656, df = 1, p-value = 0.1969 |
|  | Olfaction_Sensory_cortex | 9 | 3 | 0.07258065 | 0.06000000 | X-squared = 0.087832, df = 1, p-value = 0.767 |
|  | Olfaction_Motor | 4 | 0 | 0.03225806 | 0.00000000 | X-squared = 1.6509, df = 1, p-value = 0.1988 |
|  | Olfaction_Neuromodulators | 3 | 0 | 0.02419355 | 0.00000000 | X-squared = 1.2309, df = 1, p-value = 0.2672 |
| PREFRONTAL | PFC_all | 31 | 24 | 0.25 | 0.48 | **X-squared = 8.7194, df = 1, p-value = 0.003148 \*\*** |
|  | Intra_PFC | 4 | 7 | 0.03225806 | 0.14000000 | **X-squared = 6.9844, df = 1, p-value = 0.008222 \*\*** |
|  | Olfaction_PFC | 8 | 2 | 0.06451613 | 0.04000000 | X-squared = 0.39537, df = 1, p-value = 0.5295 |
|  | PFC_Memory | 8 | 6 | 0.06451613 | 0.12000000 | X-squared = 1.4826, df = 1, p-value = 0.2234 |
|  | PFC_Sensory_cortex | 5 | 0 | 0.04032258 | 0.00000000 | X-squared = 2.0758, df = 1, p-value = 0.1497 |
|  | PFC_Motor | 4 | 4 | 0.03225806 | 0.08000000 | X-squared = 1.8516, df = 1, p-value = 0.1736 |
|  | PFC_Neuromodulators | 2 | 5 | 0.01612903 | 0.10000000 | **X-squared = 6.4916, df = 1, p-value = 0.01084 \*** |
| MEMORY | Memory_all | 65 | 16 | 0.5241935 | 0.3200000 | **X-squared = 5.9711, df = 1, p-value = 0.01454 \*** |
|  | Intra_Memory | 13 | 1 | 0.1048387 | 0.0200000 | X-squared = 3.4664, df = 1, p-value = 0.06263 |
|  | Olfaction_Memory | 19 | 4 | 0.1532258 | 0.0800000 | X-squared = 1.6656, df = 1, p-value = 0.1969 |
|  | PFC_Memory | 8 | 6 | 0.06451613 | 0.12000000 | X-squared = 1.4826, df = 1, p-value = 0.2234 |
|  | Memory_sensory_cortex | 10 | 2 | 0.08064516 | 0.04000000 | X-squared = 0.91677, df = 1, p-value = 0.3383 |
|  | Memory_Motor | 9 | 2 | 0.07258065 | 0.04000000 | X-squared = 0.63868, df = 1, p-value = 0.4242 |
|  | Memory_Neuromodulators | 6 | 1 | 0.0483871 | 0.0200000 | X-squared = 0.74365, df = 1, p-value = 0.3885 |
| MOTOR | Motor_all | 23 | 9 | 0.1854839 | 0.1800000 | X-squared = 0.0071397, df = 1, p-value = 0.9327 |
|  | Intra_Motor | 1 | 0 | 0.008064516 | 0.000000000 | X-squared = 0.40556, df = 1, p-value = 0.5242 |
|  | Motor_Olfaction | 4 | 0 | 0.03225806 | 0.00000000 | X-squared = 1.6509, df = 1, p-value = 0.1988 |
|  | Motor_PFC | 4 | 4 | 0.03225806 | 0.08000000 | X-squared = 1.8516, df = 1, p-value = 0.1736 |
|  | Motor_Memory | 9 | 2 | 0.07258065 | 0.04000000 | X-squared = 0.63868, df = 1, p-value = 0.4242 |
|  | Motor_Sensory_cortex | 4 | 2 | 0.03225806 | 0.04000000 | X-squared = 0.064147, df = 1, p-value = 0.8001 |
|  | Motor_Neuromodulators | 1 | 1 | 0.008064516 | 0.020000000 | X-squared = 0.44675, df = 1, p-value = 0.5039 |
| SENSORY | Sensory_all | 32 | 8 | 0.2580645 | 0.1600000 | X-squared = 1.9355, df = 1, p-value = 0.1642 |
|  | Intra_Sensory | 2 | 1 | 0.01612903 | 0.02000000 | X-squared = 0.031511, df = 1, p-value = 0.8591 |
|  | Sensory_Olfaction | 9 | 3 | 0.07258065 | 0.06000000 | X-squared = 0.087832, df = 1, p-value = 0.767 |
|  | Sensory_PFC | 5 | 0 | 0.04032258 | 0.00000000 | X-squared = 2.0758, df = 1, p-value = 0.1497 |
|  | Sensory_Memory | 10 | 2 | 0.08064516 | 0.04000000 | X-squared = 0.91677, df = 1, p-value = 0.3383 |
|  | Sensory_Motor | 4 | 2 | 0.03225806 | 0.04000000 | X-squared = 0.064147, df = 1, p-value = 0.8001 |
|  | Sensory_Neuromodulators | 2 | 0 | 0.01612903 | 0.00000000 | X-squared = 0.81583, df = 1, p-value = 0.3664 |
| LC | LC_all | 0 | 3 | 0.00000000 | 0.06 | **X-squared = 7.5705, df = 1, p-value = 0.005933 \*\*** |
|  | LC_Olfaction | 0 | 0 | 0.00000000 | 0.00000000 | X-squared = NaN, df = 1, p-value = NA |
|  | LC_PFC | 0 | 3 | 0.00000000 | 0.06 | **X-squared = 7.5705, df = 1, p-value = 0.005933 \*\*** |
|  | LC_Memory | 0 | 0 | 0.00000000 | 0.00000000 | X-squared = NaN, df = 1, p-value = NA |
|  | LC_Motor | 0 | 0 | 0.00000000 | 0.00000000 | X-squared = NaN, df = 1, p-value = NA |
|  | LC_Sensory | 0 | 0 | 0.00000000 | 0.00000000 | X-squared = NaN, df = 1, p-value = NA |
|  | LC_Neuromodulators | 0 | 0 | 0.00000000 | 0.00000000 | X-squared = NaN, df = 1, p-value = NA |

**Table 9:**
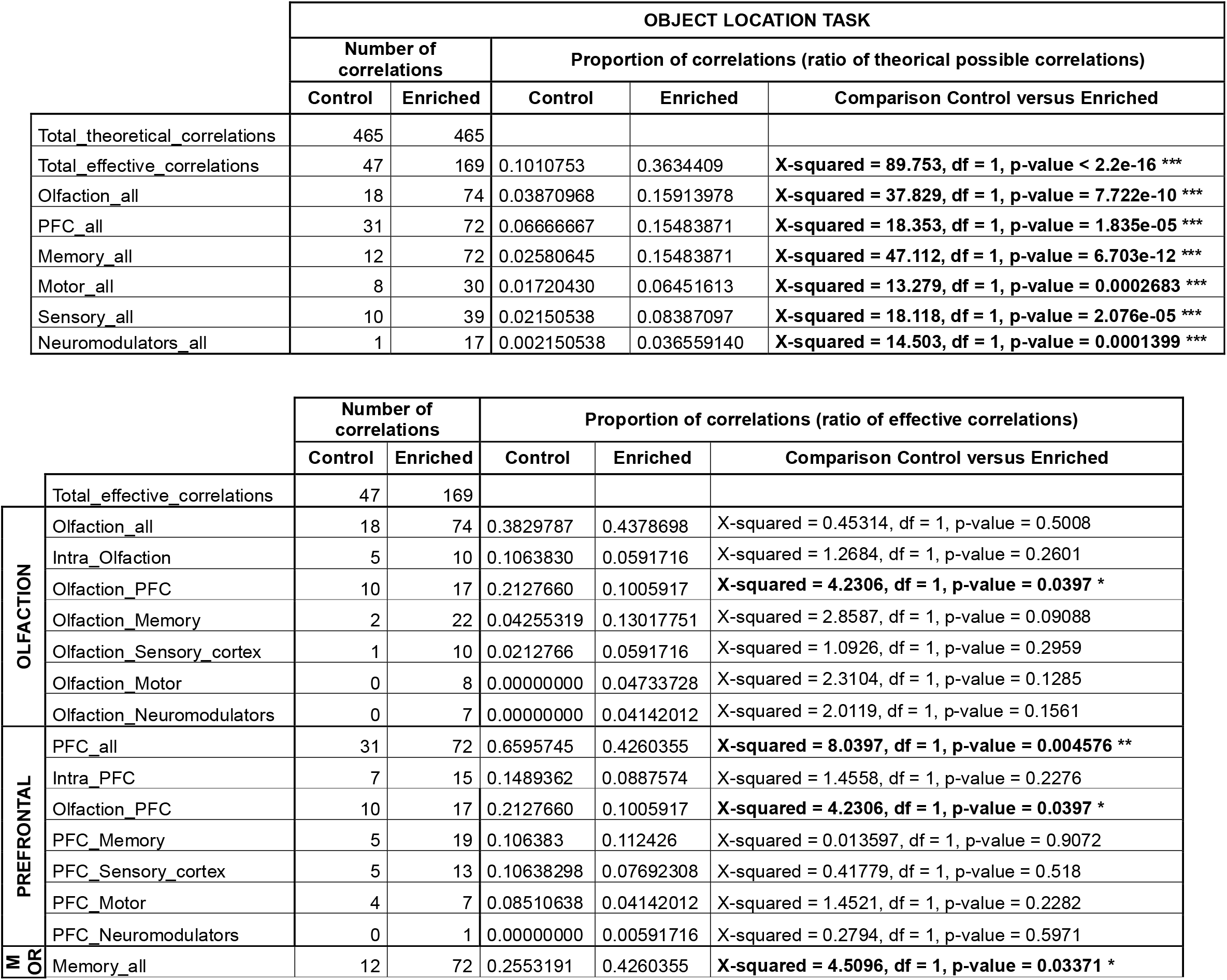
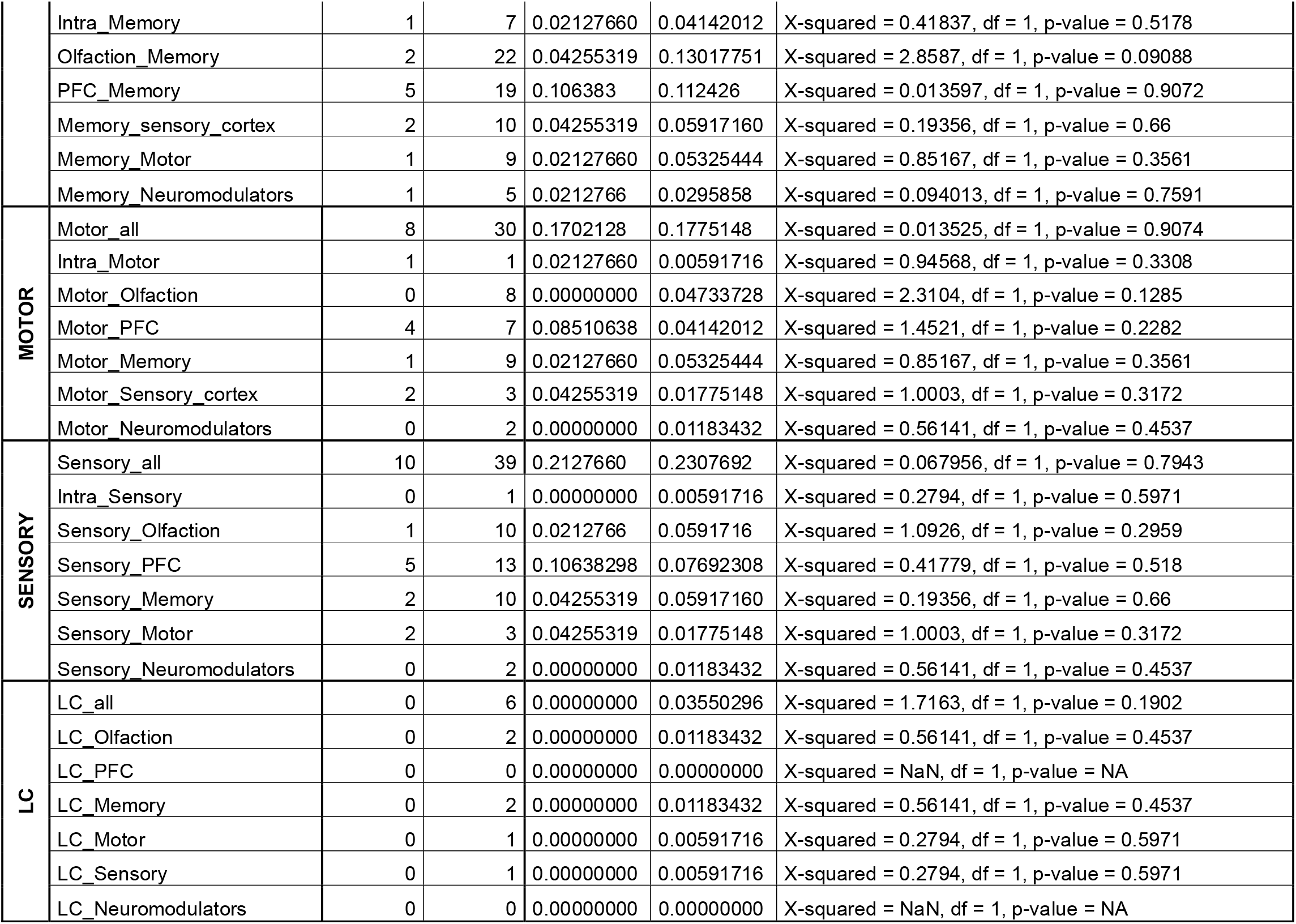
Detailed number of correlations and proportions tests for the object location task, related to Figure 8.

All together, these results indicated that a life-long olfactory enrichment induced task-dependent changes in brain functional connectivity. The olfactory discrimination task was accompanied by an increased network specialization and the spatial memory task by a network enlargement. A refinement of the network could account for the improved olfactory discrimination whereas the improved spatial memory could be driven by a more widespread brain network enabling recruitment of additional regions to perform the task.

## DISCUSSION

All together our data demonstrate benefits of long-run olfactory enrichment on age-sensitive performances that extend beyond the olfactory sphere and include olfactory discrimination, spatial memory and cognitive flexibility. These benefits correlated with a local increase of noradrenergic innervation in key olfactory and memory regions. This is accompanied by a remodeling of brain functional networks, with a more restricted and clustered network underlying efficient olfactory discrimination, while broader brain activation is associated with better spatial memory. Thus, long-run olfactory enrichment promotes plasticity of the noradrenergic system and functional network remodeling to support brain cognitive functions and appears as a pre-clinical model of the cognitive reserve buildup, opening new avenues to promote better cognitive aging.

### Long-run olfactory enrichment promotes better cognitive aging and NA structural plasticity

Here, we report that long-run repeated olfactory enrichment sessions durably improved fine olfactory discrimination, that otherwise declined with aging (our data, Enwere et al., 2004; Rey et al., 2012; Moreno et al., 2014; Yoder et al., 2017; Greco-Vuilloud et al., 2022) and was not rescued by short (30 days) olfactory enrichment (Rey et al., 2012). Noteworthily, the age-related deficits in spatial memory and cognitive flexibility were also prevented or postponed by long-run olfactory enrichment, whether started at young adulthood or at middle age. Thus, olfactory stimulation was able to induce plasticity in the middle-aged brain as observed with global environmental enrichment (*e.g.* Kobayashi et al., 2002; Harburger et al., 2007; Mora-Gallegos et al., 2015; Fuchs et al., 2016). The observation that mice with the best olfactory discrimination performances also had the best spatial memory performances, suggested that long-run olfactory enrichment activated a network that was large enough to modulate other than olfactory brain functions. In support of this hypothesis, olfactory inputs reach the hippocampus (Soudry et al., 2011) where olfactory cues are represented to notably support odor learning (Woods et al., 2020) or spatial navigation (Fischler-Ruiz et al., 2021). Interestingly, at 18 months, we showed an increase in the density of NA innervation in the olfactory bulb and dorsal CA1, correlated with olfactory discrimination and spatial memory performances respectively, suggesting a role for NA innervation plasticity in the improvement of performances. Increased NA innervation is likely to result in an increased NA release by *Locus Coeruleus* fibers. In the olfactory bulb, NA is reported to modulate in a complex way the gain of the mitral to granule cells loop (Mouly et al., 1995; Ciombor et al., 1999; Nai et al., 2009, 2010; Zimnik et al., 2013; Schwarz and Luo, 2015; Manella et al., 2017) to favor olfactory discrimination (Doucette et al., 2007; Escanilla et al., 2010; Linster, 2019). Regarding the hippocampus, NA release contributes to spatial memory by enhancing hippocampal long-term plasticity in young adult mice (Li et al., 2013). Some data reported the co-release of dopamine by *Locus Coeruleus* fibers in the hippocampus, supporting the enhancement of hippocampal memory induced by environmental enrichment (Kempadoo et al., 2016; Takeuchi et al., 2016; Wagatsuma et al., 2018). It thus cannot be excluded that dopamine contributes to the spatial memory improvements we observed in enriched aged mice.

Looking at the control groups, we observed differences in performances between early- and late-control mice at age 18 months. More precisely, late-controls were unable to learn the odor-reward association while early-controls were. Aside from a cohort effect, this difference could be due to the repeated manipulation of the animals. Control groups were manipulated as much as enriched animals and exposed to an empty (non-odorized) tea ball, that could constitute an enrichment of the living conditions (van Praag et al., 2000) and could influence their learning and memory capacities. This could as well explain the better performances in discrimination displayed by the early control group compared to the late control groups at 12 months and in spatial memory at 15 months. Importantly, these observations highlight the necessity of sham manipulations, as performed in this study, to reveal the specific effect of odor exposure versus other environmental manipulations on cognition and brain plasticity. Consequently, the effect of long-run olfactory enrichment that we described here, can thus most likely be ascribed to the repeated olfactory stimulation.

### Mechanisms underlying noradrenergic fiber structural plasticity

Based on the literature associating the anatomical integrity of the *Locus Coeruleus* with the preservation of cognitive functions in elderly (Robertson, 2013; Wilson et al., 2013; Clewett et al., 2016; Mather and Harley, 2016; Dahl et al., 2019), we hypothesized that long-run olfactory enrichment would repeatedly stimulate and thereby preserve this neuromodulatory system and favor cognitive performances during aging. In line with the broad innervation provided by the *Locus Coeruleus* to the anterior brain, we observed an increased NA innervation in the olfactory bulb and hippocampus (CA1) in response to long-run olfactory enrichment. No such plasticity was observed in the lateral orbital cortex or other hippocampal regions (CA3, DG). One possible explanation may come from the fact that the *Locus Coeruleus* is no longer considered as a homogeneous group of neurons providing a uniform signal to the brain. Indeed, there is accumulating evidence for specialization of *Locus Coeruleus* neurons into subsets displaying functional heterogeneity (patterns of discharge) and specific efferent and afferent connectivity (Chandler et al., 2014; Schwarz and Luo, 2015; Schwarz et al., 2015; Kebschull et al., 2016; Uematsu et al., 2017; Poe et al., 2020; Breton-Provencher et al., 2021). It is therefore possible that long-run olfactory enrichment stimulated a subset of *Locus Coeruleus* neurons.

The NA structural plasticity that we observed could be due to sprouting of axon collaterals of *Locus Coeruleus* neurons within their target regions since the number of NA neurons in the *Locus Coeruleus* remained similar in control and enriched groups. Such remodeling of NA fibers had not been shown in the olfactory bulb following olfactory enrichment of shorter duration (Rey et al., 2012), suggesting that the repetition of the enrichment period is necessary to trigger such plasticity. Remodeling of NA fibers is well described after lesioning of the *Locus Coeruleus* (*e.g.* Fritschy and Grzanna, 1992) or of its projections (*e.g.* Peterson, 1994) but only one study reported NA fiber remodeling in physiological conditions within the cerebellum following exercise (Nedelescu et al., 2017). Thus, our study provides important evidence for physiological structural plasticity of the NA innervation within cortical regions and for the occurrence of this process in aged mice.

### Brain networks remodeling induced by long-run olfactory enrichment and its contribution to olfactory and cognitive improvements

The behavioral improvements observed in aged mice after long-run olfactory enrichment were accompanied by a remodeling of brain activation patterns. Importantly, this remodeling emerged from the comparison of two groups of aged animals (control and enriched) performing the task but only one, the enriched group, performed it successfully. This suggests that this remodeling enables performance improvement after long-run olfactory enrichment. A further argument supporting a role of network remodeling in such improvement arises from the observation that this remodeling was task-specific. More precisely, an increased network specialization was associated with the olfactory discrimination task (less connections and more clusters) and a network enlargement with the spatial memory task (more connections and more regions participating to the network). Remarkably, in the olfactory discrimination task, the relative contribution of connections within the olfactory system to the whole network increased, suggesting a specialization of the network in regard to the olfactory task after long-run olfactory enrichment. A remodeling consistent with these observations has been reported in anosmic patients submitted to olfactory enrichment, who displayed improved detection thresholds and increased number of functional connections in the odor-evoked network (Kollndorfer et al., 2015). Furthermore, these connections were more specific as the piriform cortex was connected to fewer non-olfactory structures (Kollndorfer et al., 2014). In addition, the relative contribution of the connectivity within PFC structures and of that between PFC structures and the neuromodulatory systems increased in the olfactory discrimination task, possibly reflecting a reinforcement of attention processes after long-run olfactory enrichment. In the object location task, a denser and larger network was activated in enriched animals with more connections and the recruitment of new regions. Making sense with the spatial memory behavioral task, some of these regions are strongly involved in spatial memory (*e.g.* Moser et al., 1995, 2017; Assini et al., 2009; Fanselow and Dong, 2010; Eichenbaum, 2017; Igarashi et al., 2022) (entorhinal cortex and dCA1). The recruitment of additional brain regions has also been observed in aged rats succeeding in an aversive olfactory associative task, when compared to young animals (Dardou et al., 2010). Altogether, this supports the view that the network enlargement reported here could support the spatial memory in aged enriched mice.

More generally, lifelong olfactory enrichment promotes cognition and large-scale and task-dependent network changes. This is reminiscent of what has been observed in aged humans in which rearrangements of brain connectivity were described and linked to performances. These results suggest that common mechanisms of aged-related network remodeling could be at play in humans and animals. In humans, both specification and enlargement of network reorganization can underly preserved cognitive functions (Grady, 2012; Cabeza et al., 2018; Crowell et al., 2020). Depending on the task, additional recruitment of brain areas in elderly can either be accompanied by an improvement (Fera, 2005; Crowell et al., 2020) or a decrease in performance (Persson and Nyberg, 2006). In the latter case, the increase in activity observed in some regions would not reflect compensation but rather a lack of specificity or less efficient activity (Morcom and Henson, 2018). Our data suggest that following long-run olfactory enrichment, the overall increased connectivity is associated with better performances in spatial memory in aged mice.

Last, *Locus Coeruleus* activation and NA sprouting could contribute to the observed behavioral improvement by remodeling brain activation networks. In support to this view, we observed that the *Locus Coeruleus* became more connected with other brain regions in enriched animals, for both olfactory discrimination and spatial memory tasks. This suggests that long-run olfactory enrichment, in line with our hypothesis, favors the maintenance of the functional connectivity of the *Locus Coeruleus* and thereby the supply of NA to cortical regions. A causal involvement of NA structural plasticity in the network remodeling and in the improvement of the performances in the aged enriched animals remains to be demonstrated but is supported by the report of a functional reconfiguration of brain neural networks (activation and connectivity) induced by the chemogenetic activation of the NA system in the young mouse brain (Zerbi et al., 2019).

In conclusion, this study revealed long-run olfactory enrichment as an efficient strategy to broadly improve cognition in aged mice, by inducing localized increase of NA innervation and brain network remodeling. We thus propose that the long-run olfactory enrichment in mice should be considered as an interesting model of cognitive reserve buildup, that could contribute to the identification of underlying cellular and sub-cellular mechanisms, that cannot be explored in humans. In combination with other clinical interventions, this non-invasive non-pharmaceutical remediation strategy seems also promising to promote better brain aging in humans for at least two reasons. First, it is based on a single sensory modality and would be easy to transpose at low cost, compared to global environmental enrichment, by proposing odor sets or odor-based games as leisure activities. Second, throughout the literature in animals, the efficiency for improving cognition with environmental or strictly olfactory enrichment, relies on novelty exposure (van Praag et al., 2000; Veyrac et al., 2009; Li et al., 2013). The olfactory environment of most humans, if not poor in the sense that everyone is exposed to various odors every day, is likely to be rather stable, composed mostly of familiar odorants. Thus, there is a real scope for increasing the variety of smells to which we are exposed and this could be relevant in real life, and not only in laboratory animals, to trigger the mechanisms of the cognitive reserve build up. Finally, adding to the translational value of olfactory enrichment, late-started olfactory enrichment improves spatial memory as efficiently as early-started enrichment, indicating that the olfactory strategy of promoting better aging may be efficient even when started late in life.

## ACKNOWLEDGEMENT

This work was supported by the CNRS, Inserm, Lyon 1 University, a PhD fellowship from the Arc2 Région Auvergne Rhône Alpes to C.T, a PhD fellowship from Fondation de France – Fondation Roudnitska to J.G-V and a PhD fellowship from Ecole Normale Supérieure de Lyon to M.C. We thank Ounsa Ben Hellal-Jelassi for her valuable care to the aging mouse colony.

## SUPPLEMENTARY FIGURES AND LEGENDS

**Figure S1:**
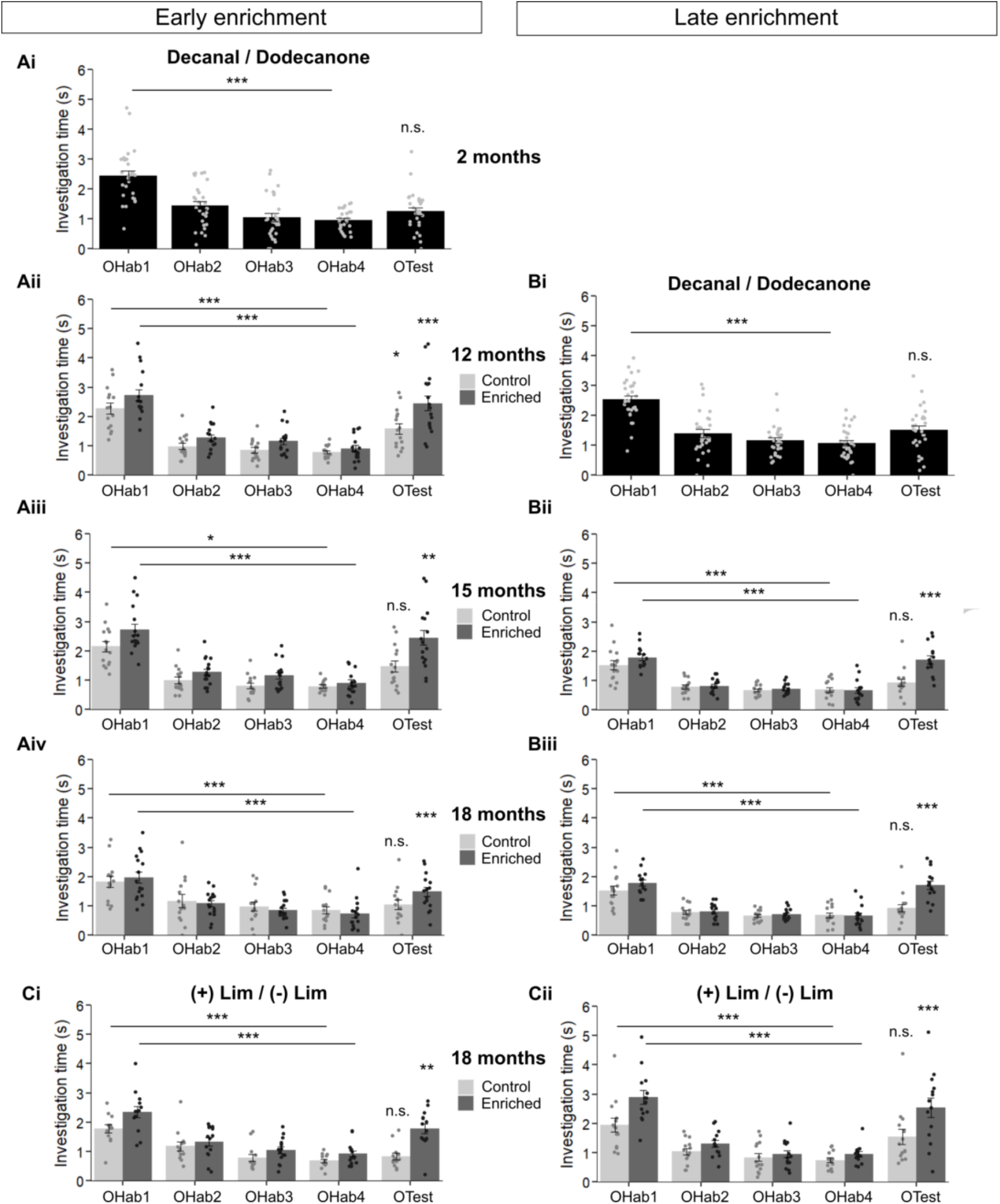
Habituation to odorants and discrimination of perceptually similar odorants after long-run olfactory enrichment. Assessment of fine olfactory discrimination at different ages across life-time in the enriched and control groups of early- and late-started enrichment, using a cross habituation task with perceptually similar odorant pairs Decanal / Dodecanone **(A-B)** or (+) Limonene / (-) Limonene **(C)**. All groups of mice habituated to the first presented odorant (OHab) for both odorant pairs, as evidenced by the significantly decreased investigation time between OHab1 and OHab4. Discrimination was tested by comparing the investigation time between OHab4 and OTest. In pre-enrichment tests, mice did not discriminate Decanal / Dodecanone (**Ai and Bi**, p>0.05). Both early- and late-enriched mice were able to discriminate Decanal / Dodecanone **(Aii-Aiv and Bii-Biii)** and (+) Limonene / (-) Limonene **(Ci-Cii)**, as shown by the significantly increased investigation time between OHab4 and OTest, while controls were not. See Fig.2 for complete description of the discrimination results. All results are given as mean ± SEM, dots represent individual mice. n= 12-28 mice per group (see Table 2 for details). * p<0.05; ** p<0.01; *** p<0.001

**Figure S2:**
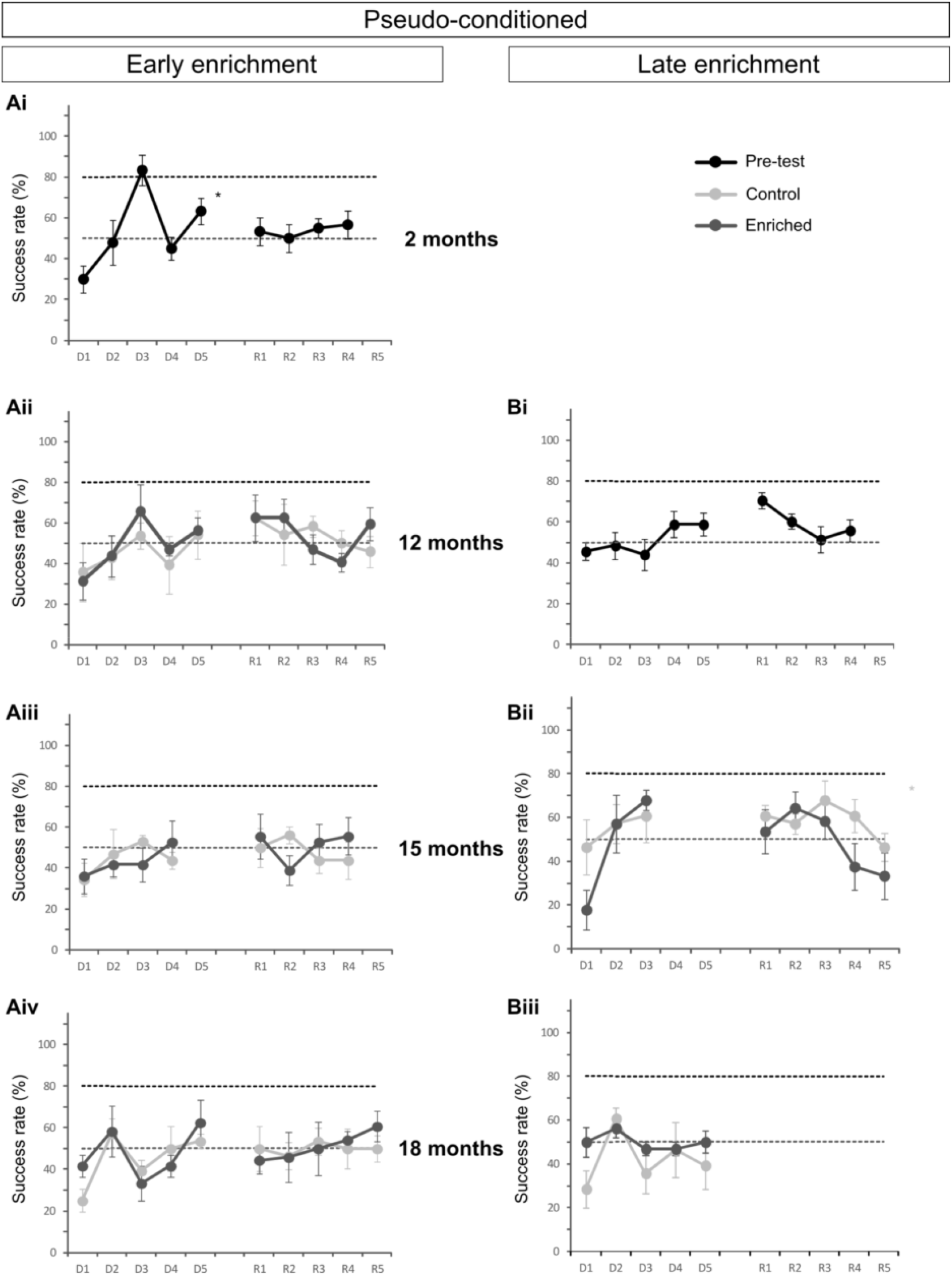
Pseudo-conditioned mice are unable to learn the olfactory associative task. Evolution of the success rate in the olfactory associative task during the acquisition of the first rule (D1-D5) and after the reversal of the rule (R1-R5) for the pseudo-conditioned groups. Mice were tested before the beginning of the enrichment (pre-tests) and at different ages (12, 15 and 18 months) in the enriched and control groups of early-**(A)** and late-started **(B)** enrichment. Data are presented as mean success rate ± standard error of the mean. n=6-17 mice per group (details are provided in Table S1).

**Figure S3:**
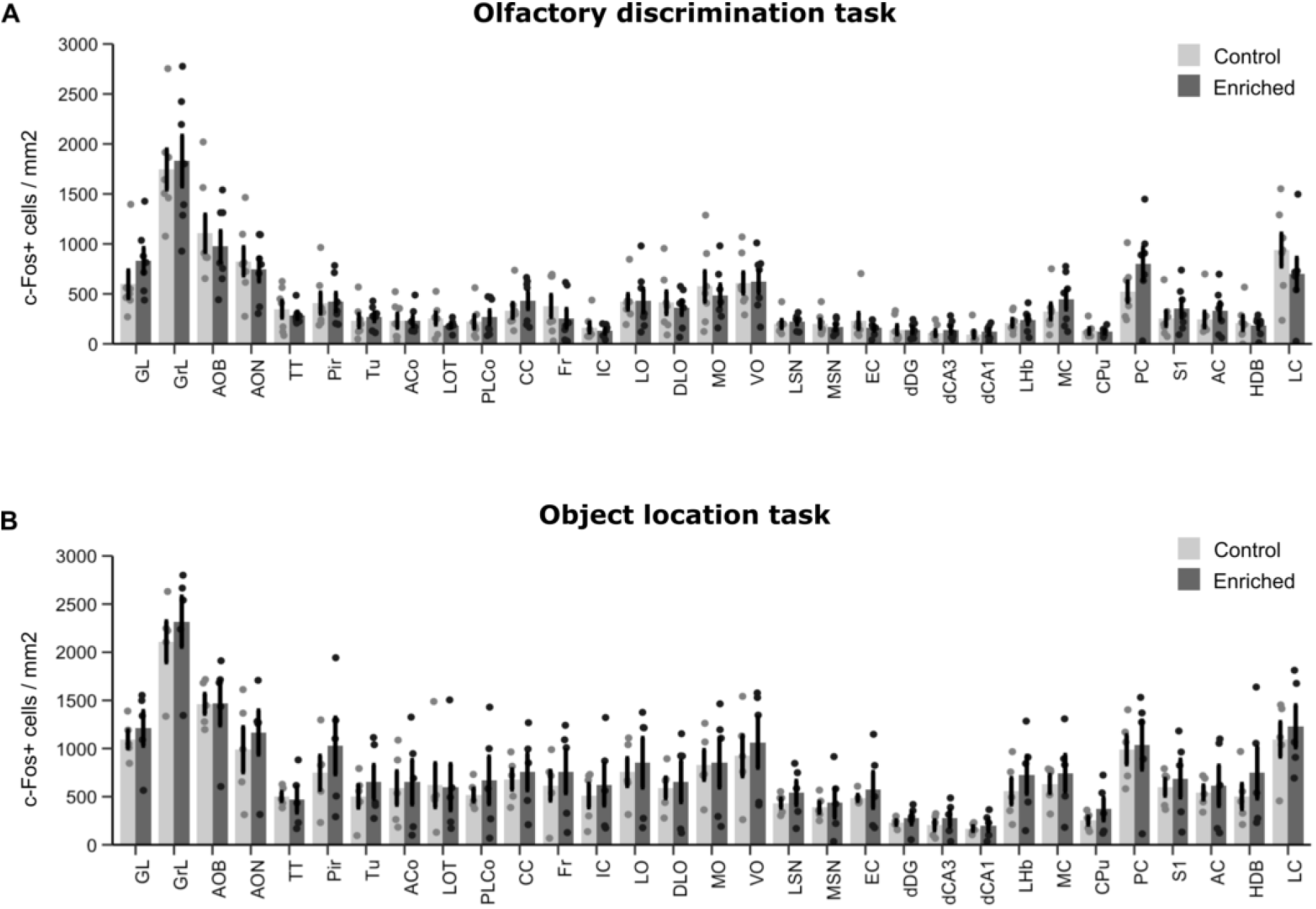
Brain regional activation induced by olfactory discrimination task and object location task (raw data before normalization as in Figure 7). c-Fos+ cell densities of enriched mice for the 31 regions of interest in the olfactory discrimination task **(A)** and the object location task **(B)**. Data represent the mean ± SEM. Dots represent data from individual mice. n=5-7 mice per group. For the olfactory discrimination task, data normality hypothesis was not verified. Comparisons of control vs enriched for each structure by bilateral unpaired Wilcoxon test with Holm’s correction for multiple comparisons yielded no differences. After pooling control and enriched groups, Kruskal-Wallis test identified a significant effect of structure (H (30) = 230.3372, p < 0.001). For the object location task, data homoscedasticity hypothesis was not verified. Comparisons of control vs enriched for each structure by Student t-test with Welch correction and Holm’s correction for multiple comparisons yielded no differences. After pooling control and enriched groups, Kruskal-Wallis test identified a significant effect of structure (H (30) = 236.7178, p < 0.001).

**Figure S4:**
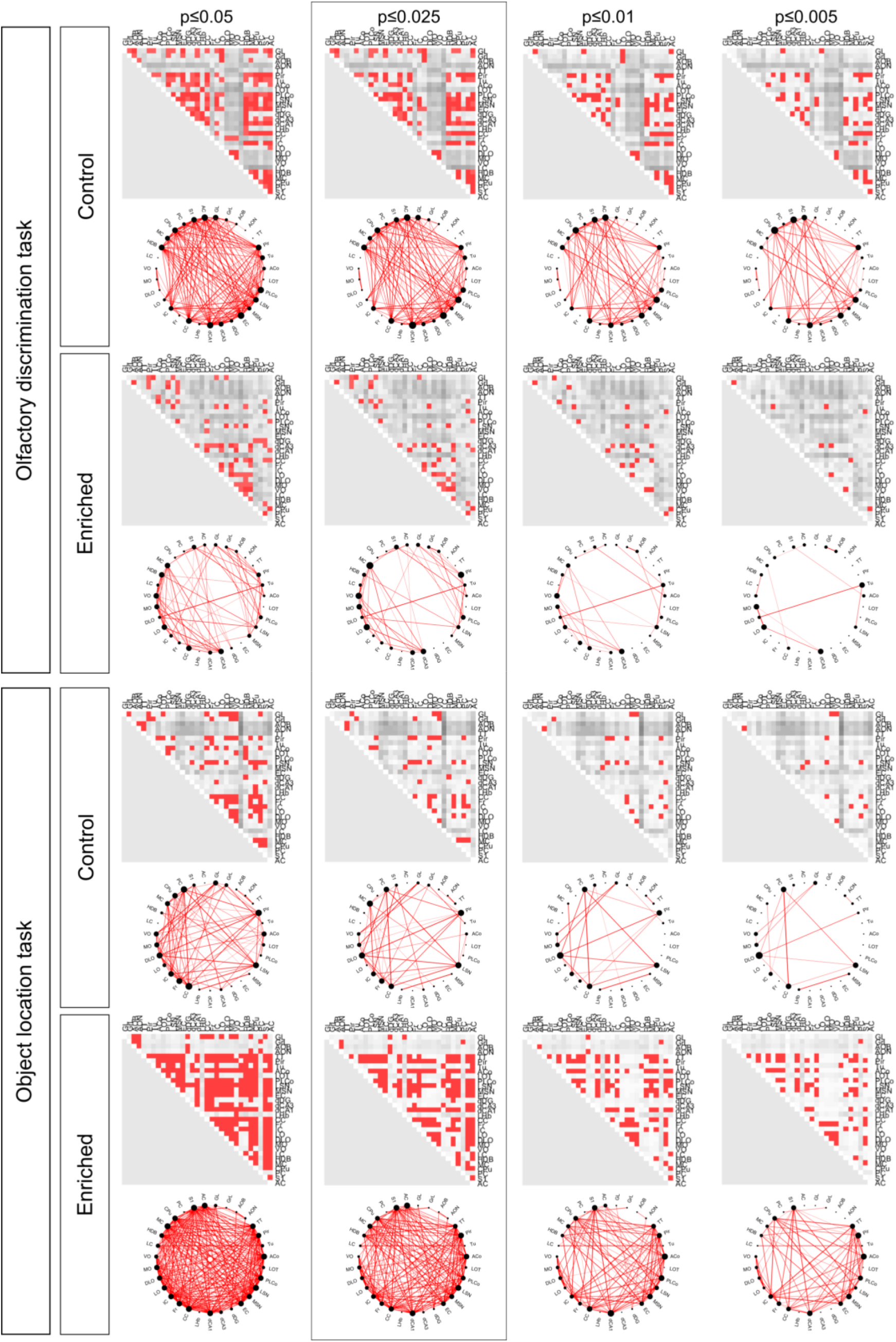
Correlations matrices obtained with different threshold levels reveal comparable differences between control and enriched groups. Matrices representing all inter-regional correlations with threshold p-values of p≤0.05, p≤0.025, p≤0.01 and p≤0.005. Axes correspond to brain structures and red color reflects correlations above threshold. p≤0.025 was the selected p-value for the data presented in Fig8.

**Figure S5:**
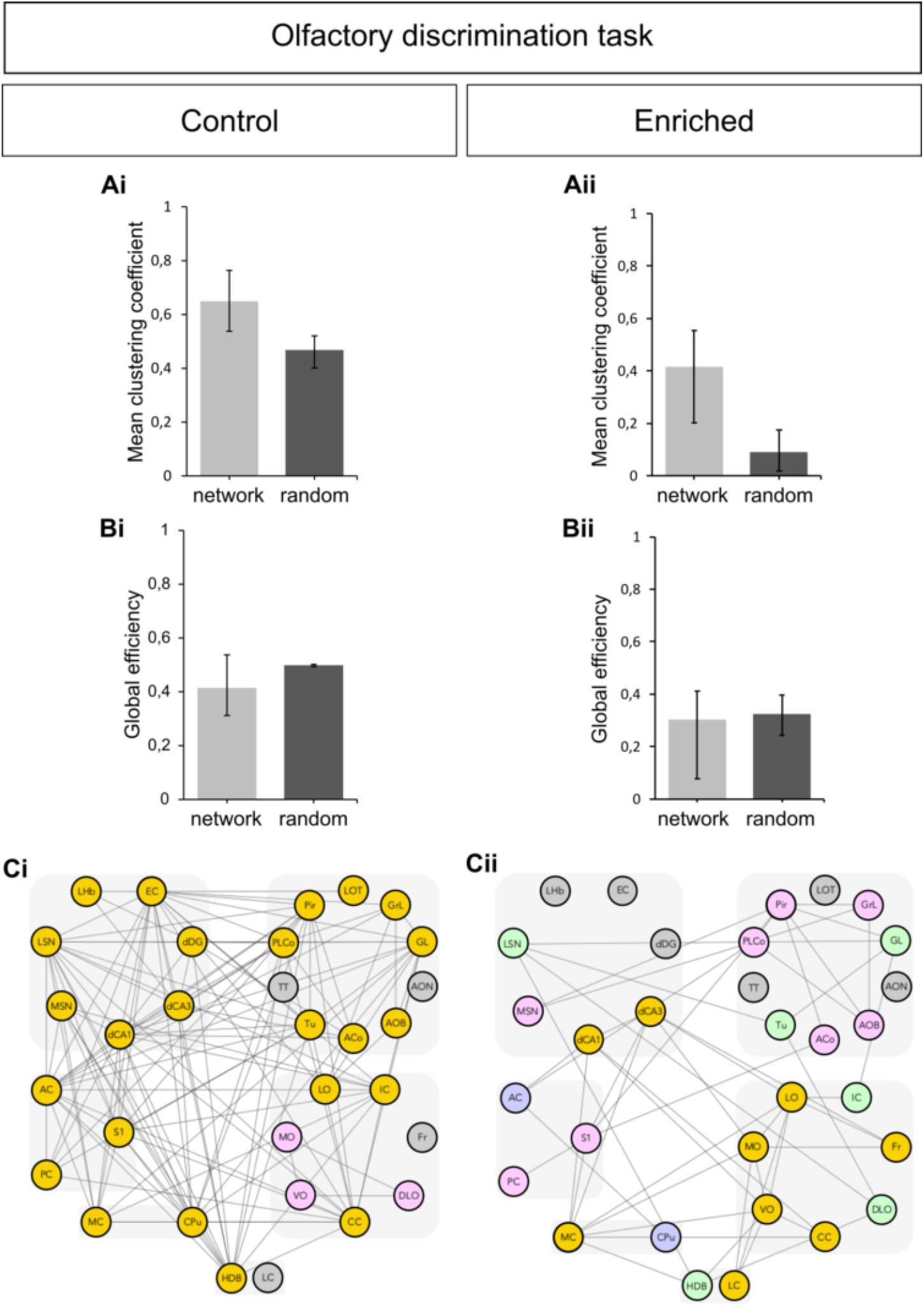
The olfactory discrimination network displays a small-world organization. **(A)** Mean clustering coefficient was measured to assess the segregation of the networks for control **(Ai)** and enriched **(Aii)** groups that performed the olfactory discrimination task. **(Bi-Bii)** Global efficiency was measured to assess the integration of the networks. Each parameter is compared between the experimental network and a random network. Error bars represent 95% confidence intervals. **(Ci-Cii)** Other representation of the networks organized into discrete color-coded clusters (pink, green, blue, yellow) based on their common inter-connections using the Markov clustering algorithm. Regions that are not correlated at all are represented in grey. Two clusters were identified in the control group **(Ci)**: a small cluster involving prefrontal areas and a widespread cluster including olfactory, spatial memory, motor and sensory areas. Four clusters were found in the enriched group **(Cii)**: a first one involving mostly olfactory regions (pink); a second involving the reward system (green); a third involving auditory and motor structures (blue); and the fourth (yellow) involving prefrontal structures, spatial memory structures, motor structures and the *Locus Coeruleus*.

**Figure S6:**
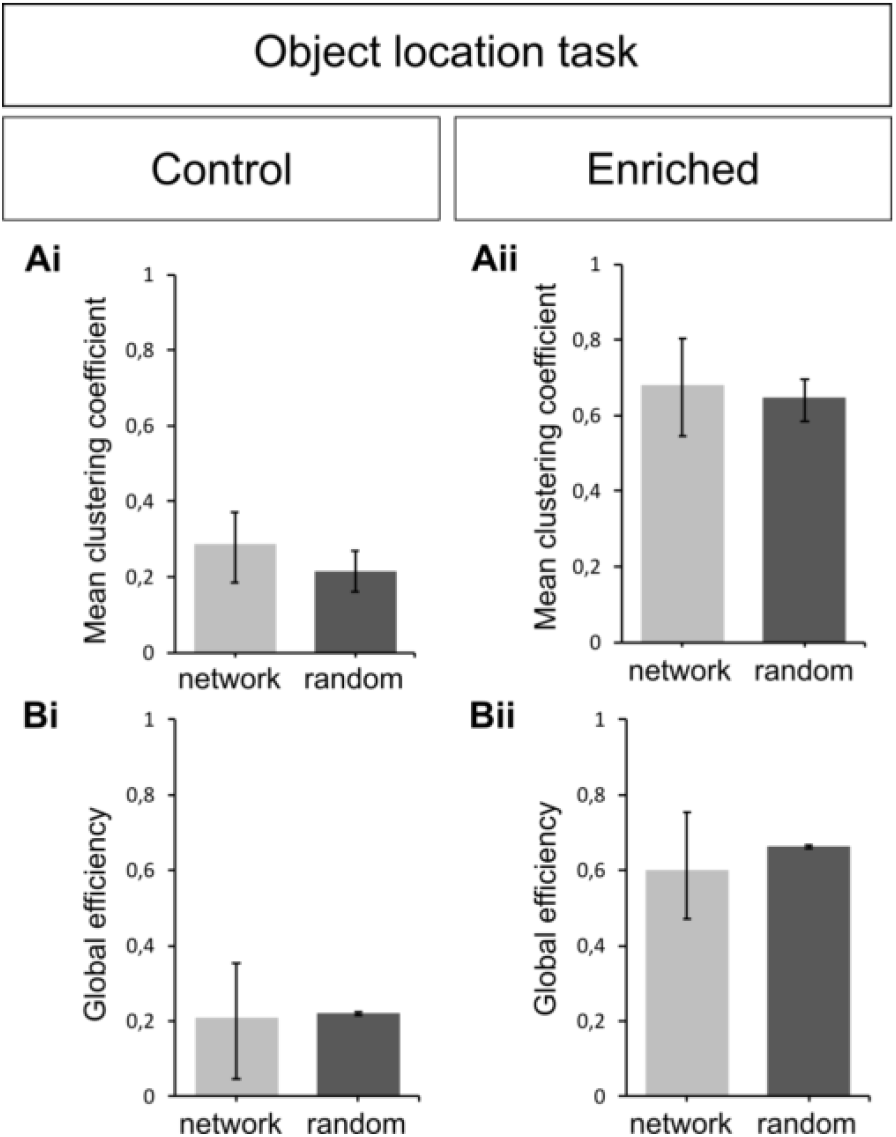
The object location network does not display a small-world organization. **(Ai-Aii)** Mean clustering coefficient was measured to assess the segregation of the networks for control and enriched groups that performed the object location task. **(Bi-Bii)** Global efficiency was measured to assess the integration of the networks. Each parameter is compared between the experimental network and a random network. Error bars represent 95% confidence intervals.

**Table S1:**
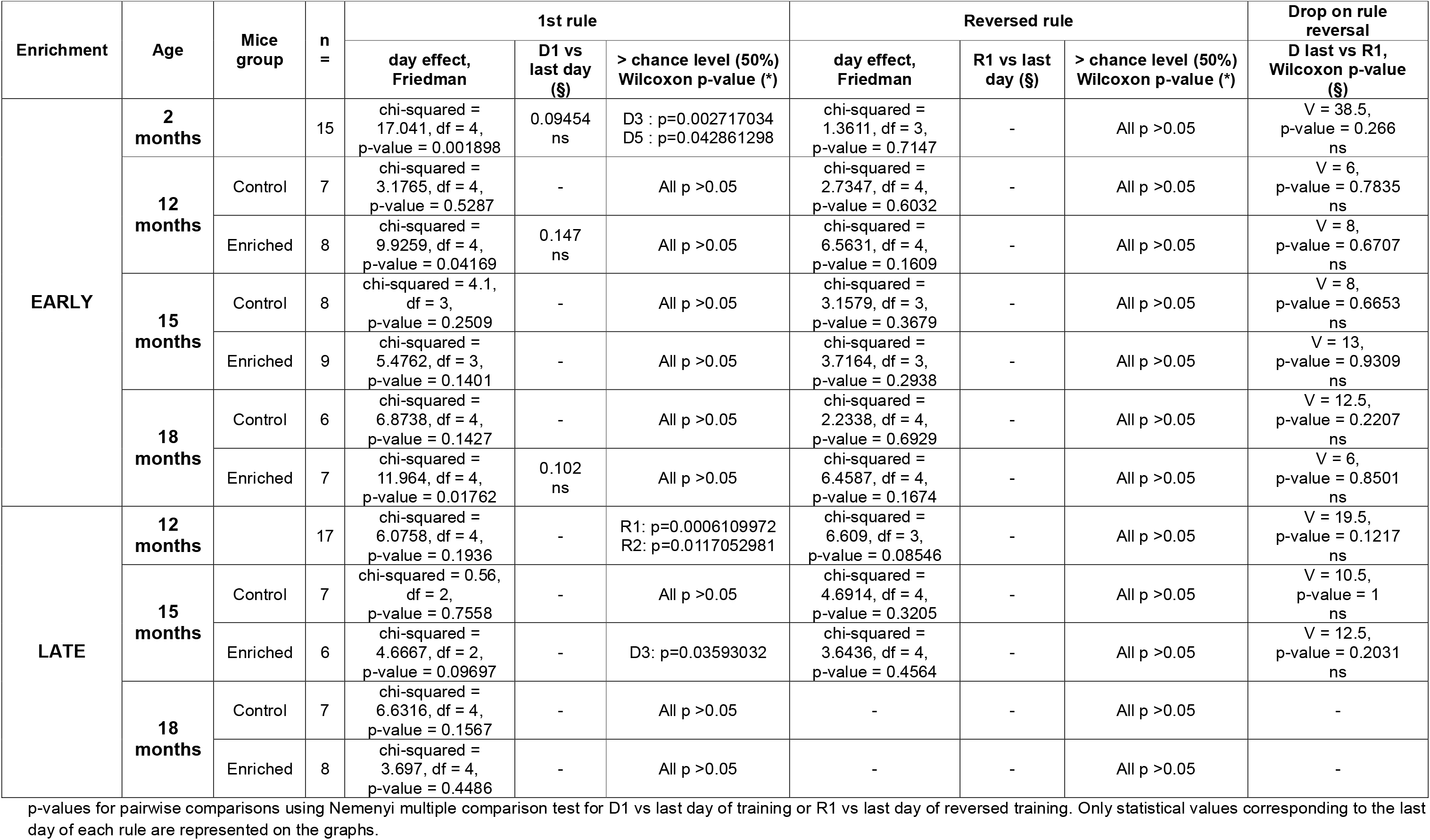
Detailed statistical analysis related to Figure S2.

